# Proteomic Investigation of Neural Stem Cell to Oligodendrocyte Precursor Cell Differentiation Reveals Phosphorylation-Dependent Dclk1 Processing

**DOI:** 10.1101/2023.04.17.537129

**Authors:** Robert Hardt, Alireza Dehghani, Carmen Schoor, Markus Gödderz, Nur Cengiz Winter, Shiva Ahmadi, Ramesh Sharma, Karin Schork, Martin Eisenacher, Volkmar Gieselmann, Dominic Winter

## Abstract

Oligodendrocytes are generated via a two-step mechanism from pluripotent neural stem cells (NSCs): after differentiation of NSCs to oligodendrocyte precursor/NG2 cells (OPCs), they further develop into mature oligodendrocytes. The first step of this differentiation process is only incompletely understood. In this study, we utilized the neurosphere assay to investigate NSC to OPC differentiation in a time course-dependent manner by mass spectrometry-based (phospho-) proteomics. We identify double cortin like kinase 1 (Dclk1) as one of the most prominently regulated proteins in both datasets, and show that it undergoes a gradual transition between its short/long isoform during NSC to OPC differentiation. This is regulated by phosphorylation of its SP-rich region, resulting in inhibition of proteolytic Dclk1 long cleavage, and therefore Dclk1 short generation. Through interactome analyses of different Dclk1 isoforms by proximity biotinylation, we characterize their individual putative interaction partners and substrates.

## Introduction

The central nervous system (CNS) contains a plethora of cell types which are commonly grouped into two major categories, neurons and glia, each accounting for ∼50% of cells.^1^ Neurons are responsible for signal transduction and processing of information, and many different sub-types are known today, which can be distinguished by their function and/or protein expression patterns, even within individual structures.^2^ Glia provide multiple other functions for the central nervous system and are commonly further divided into four main classes. Astrocytes support neurons, maintaining metabolic and structural homeostasis; microglia function as immune cells of the CNS; oligodendrocytes (ODs) form myelin sheaths surrounding the axons of neurons, improving signal transmission and promoting neuronal repair and development;^1^ and NG2 cells – or oligodendrocyte precursor cells (OPCs) – are mainly known for their ability to differentiate into myelinating ODs.^3^ It is estimated that OPCs contribute 5% to 8% of total glial cells in the mature CNS.^4^ While the majority of white matter OPCs can differentiate into ODs during an organism’s life span, most gray matter OPCs retain their phenotype during postnatal life, implying other functions than acting as precursors for ODs.^5^

Loss or malfunction of ODs leads to severe diseases, which can be classified in such with a genetic background, the so-called leukodystrophies, and such induced by trauma or autoimmune reactions, with multiple sclerosis being the most prominent example.^6, 7^ The common phenotype of these disorders is the inability of formation, or the progressive loss, of myelin sheaths, leading to impaired neuronal signal transduction with severe consequences for the affected individual.^8^ In a healthy organism, the loss of ODs, and the resulting demyelination, can be compensated by proliferation of OPCs and their differentiation to ODs, resulting in remyelination of axons. In demyelinating disorders, however, this ability is impaired, and OPCs, even though being present at the site of demyelination, do not differentiate into ODs.^9^

OPC to OD differentiation is one out of two steps in the transformation from pluripotent neural stem cells (NSCs) to ODs: initially, NSCs differentiate into OPCs, which subsequently further differentiate into myelinating ODs.^10^ A better understanding of these differentiation processes holds the promise to identify targets for therapeutic intervention with respect to the generation of mature ODs, ultimately facilitating increased rates of remyelination. This results in a broad interest for the underlying mechanisms. The differentiation of OPCs to ODs has been investigated in several transcriptomics/proteomics studies, which is partly also due to the fact that this process can be reproduced *in vitro* (summarized by Schoor et al).^11^ In contrast to OPC to OD differentiation, NSC to OPC differentiation has not been thoroughly investigated by unbiased screening approaches so far, including, to our knowledge, just one proteomic analysis which compared the expression levels of ∼3,000 proteins for *in vitro* cultivated human embryonic stem cells (ESCs), NSCs, OPCs, and glial progenitor cells.^12^ While these experiments were performed with ESCs cultivated on a feeder layer of mouse embryonic fibroblasts (MEFs), two other studies investigated microRNA and gene expression during NSC to OPC differentiation, respectively, utilizing so-called neurospheres.^13, 14^

Neurospheres are free-floating clusters of cells containing NSCs. They are typically generated from mouse or rat cortical tissue-derived cells which, after supplementation of the culture medium with bFGF and EGF, give rise to neurospheres that can be maintained in a progenitor state under cell culture conditions.^15^ In neurospheres, growth factor withdrawal triggers differentiation into a mixture of different CNS cell types (astrocytes, neurons, and OPCs),^16^ while treatment with cell culture medium conditioned by the neuroblastoma cell line B104 leads to the formation of so-called oligospheres, which predominantly consist of OPCs.^15^ As this allows for the generation of large numbers of OPCs in vitro, this practice is a common approach to produce OPCs, especially from mice, as the enrichment and cultivation of primary mouse OPCs is difficult.^15^

Neurosphere to oligosphere differentiation allows to investigate the differentiation of NSCs to OPCs *in vitro*. With respect to whole proteome analyses, this system has the advantage that NSCs can be cultivated without the need of a feeder layer and that a gradual increase of OPCs is observed.^15^ Therefore, no proteins/mRNAs of feeder cells contaminate the sample, and the differentiation process can be investigated in a time-course dependent manner, allowing for the identification of co-regulated proteins based on their abundance profiles.^17^ Naturally, this cell culture model cannot fully reflect the *in vivo* conditions. It was shown, however, that the expression of marker proteins during NSC to OPC differentiation *in vitro* reflects their sequence *in vivo;* demonstrating that significant parts of the differentiation program are intrinsic to these cells, and confirming the validity of studying this process *in vitro*.^18^

In the current study, we investigated the differentiation of neurospheres to oligospheres by quantitative mass spectrometry-based proteomics and phosphoproteomics. Among others, we identified the protein double cortin like kinase 1 (Dclk1) to be highly regulated both on the protein and phosphorylation site level. Further investigation revealed that differential proteolytic processing of Dclk1 correlates with phosphorylation in its SP rich region, resulting in different proteolytically processed isoforms, for which we investigated putative interaction partners and substrates using proximity biotinylation.

## Materials and Methods

### Neurosphere generation and differentiation

Breeding and handling of mice, as well as harvesting of tissues, was performed in accordance with the policies regarding animal handling and welfare of the state of North Rhine Westphalia, Germany. Neurospheres were generated as described elsewhere with some modifications.^15^ Briefly, P0/P1 mice were decapitated, cortices isolated, and placed in ice-cold HBSS. After removal of meninges, tissues were chopped into small pieces with a razor blade and placed into ice-cold Neurosphere Growth Medium (NGM, DMEM/F12 with 25 μg/ml insulin, 100 μg/ml Apo-transferrin, 20 nM progesterone, 60 μM putrescine, 30 nM sodium selenite, 100 IU/ml penicillin, 100 IU/ml streptomycin, 2 mM L-glutamine, 20 ng/ml bFGF, and 20 ng/ml EGF). For each brain, 0.5 mL of NGM was used. Cortex pieces were further dissociated into single cell suspensions using fire-polished Pasteur pipettes, passed through a 40 µm cell strainer, and counted using a hemocytometer. In each well of a six-well plate, 4 ml of NGM were combined with 2×10^5^ cells, and plates were incubated at 37° C with 5% CO_2_. Half of the medium was replaced with fresh NGM every other day. After ten days, neurospheres started to form, and after 14 days the differentiation process was induced using B104 neuroblastoma conditioned medium (B104 CM). For generation of B104 CM, B104 neuroblastoma cells were grown in DMEM/F12 supplemented with 10% FCS, 100 IU/ml penicillin, 100 IU/ml streptomycin, and 2 mM L-glutamine) to full confluency. Subsequently, the cells were washed with Puck’s balanced salt solution (8 g/l NaCl, 0.4 g/l KCl, 0.09 g/l Na_2_HPO_4_·7H_2_O, and 1 g/l glucose), and N2 medium (DMEM/F12 supplemented with 1x N2 supplement, Thermo Fisher Scientific, Waltham, MA) was added to the cells. After four days the medium was collected, 1 μg/ml of phenylmethylsulfonyl fluoride (PMSF) was added, and the solution centrifuged for 30 min at 2,000 g, 4° C. The supernatant was filtered using a 0.22 μm sterile filter and divided into aliquots which were stored at -80° C until further use. For induction of differentiation, neurospheres were transferred to a fresh six-well plate containing 4.5 ml of NGM and 1.5 ml of B104 CM. For a duration of two weeks, 25% of the medium was replaced with B104 CM every other day. At the individual time points, neurospheres were harvested by centrifugation for 5 min at 50 g, 4° C.

### Immunofluorescence Microscopy

Neurospheres were enzymatically and mechanically dissociated using accutase and fire polished Pasteur pipettes, and 31110^5^ cells/well were seeded on poly-L-ornithine-coated coverslips located in 24 well plates. Cells were incubated for 12 h either in NGM (neurospheres harvested at day 0) or NGM/B104 CM (neurospheres harvested at day 3, 6, and 9) containing 20 µM forskolin. Subsequently, coverslips were washed three times with 1x phosphate buffered saline (PBS) and fixed with 4% paraformaldehyde in 1x PBS, followed by permeabilization using 0.2% Triton X-100 (except for Ng2-staining). Coverslips were blocked using 2% normal goat serum and stained using the following primary antibodies in blocking solution overnight: anti-Ng2 (1:100, rabbit pAb, AB5320 Merck Millipore, Burlington, MA), anti-beta III tubulin (1:100, mouse mAb, T-1315, Dianova, Hamburg, Germany), anti-Gfap (1:100, mouse mAb, AB10062, Abcam, Cambridge, UK), anti-F4/80 (1:100, rat mAb, American Type Culture Collection, LGC Promochem, Teddington, UK), anti-Nestin (1:100, mouse mAb, 556309, BD Pharmingen, San Diego, CA), and anti-MBP (1:100, rat mAb, ab7349, Abcam). The next day, coverslips were washed with 1x TBS followed by incubation with DAPI (4,6-diamidino-2-phenylindol) and the respective secondary antibodies in blocking solution for 90 min at RT in the dark: anti-rabbit-Cy3 (1:600, goat, 111-165-144, Dianova), anti-mouse-Alexa488 (1:600, goat, A11017, Invitrogen, Carlsbad, CA), anti-mouse-Alexa546 (1:400, goat, A11018, Invitrogen), and anti-rat-Dylight488 (1:800, goat, 112-485-167, Dianova). After incubation, cover slips were washed with 1x TBS, water, and ethanol followed by a drying step at RT for 10 min. Subsequently, they were mounted with Prolong Gold (Thermo Fisher Scientific) on specimen slides, incubated over night at 4° C, and images were taken with an Axiovert 100M (Zeiss, Oberkochen, Germany) equipped with an AxioCamHR camera. Exposure times were adjusted based on secondary antibody controls. All images were recorded using the Plan Apochromat 20x/0.8 objective with the filter sets 01, 10, and 15, and analyzed using AxioVision SE64 (Zeiss).

### Cell culture, cloning, and site-directed mutagenesis

All cells were cultured at 37° C with 5% CO_2_. HEK293, HeLa, NIH/3T3, and NIH/3T3 Tet-On 3G (Takara Bio, Kusatsu, Japan) cells were cultured in DMEM, C6 glioblastoma cells in DMEM/F12, and B35 neuroblastoma cells in DMEM-GlutaMAX (Thermo Fisher Scientific). All media were supplemented with 10% FCS, 100 IU/ml penicillin, 100 IU/ml streptomycin, and 2 mM L-glutamine (with the exception of DMEM-GlutaMAX). For NIH/3T3 Tet-On 3G cells, the growth medium furthermore contained G418 (100 µg/ml) and protein expression was induced using doxycycline (0.05 μg/ml). Fetal neural stem (fns) cells^19^ were cultured in NS-A medium (Euroclone, Milan, Italy) containing 10% DMEM/F12, 1x N2 supplement, 100 IU/ml penicillin, and 100 IU/ml streptomycin. Oli-neu cells^20^ were cultivated in DMEM supplemented with 1x N2 supplement, 5 mg/L Insulin, 0.5 µM triiodothyronine, 0.5 µM L-thyroxine, 1% horse serum, 100 IU/ml penicillin, 100 IU/ml streptomycin, 2 mM L-glutamine, and 10 mM HEPES pH 7.4. For SILAC experiments, cells were cultured in SILAC DMEM (Thermo Fisher Scientific) supplemented with 10% dialyzed FCS, 100 IU/ml penicillin, 100 IU/ml streptomycin, and either unlabeled or heavy labeled lysine/arginine (Lys: ^13^C_6_^15^N_2_, Arg: ^13^C_6_^15^N_4_) for at least 5 passages. The Dclk1 gene was amplified by PCR from a cDNA library generated from P1 mouse whole brain extracts and cloned either under control of TRE3G promoter (for primers see Table S14). Site directed mutagenesis was performed as described elsewhere.^21^ Briefly, the melting temperature of mutation primers was designed to be > 80° C and PCR products were digested by DpnI for 4 h at 37° C, followed by transformation of bacteria and midi prep purification of plasmids. Cells were transfected using TurboFect transfection reagent (Thermo Fisher Scientific) according to the manufacturer’s instructions. After 24 h, medium was replaced with doxycycline-containing medium (0.05 µg/µl). Cells were incubated another 24 h and then either harvested or, after medium exchange, grown for an additional day.

For BioID experiments, fusion constructs of different DCLK1 domains and BirA* were generated by PCR and cloned into a vector under control of an RSV promotor (for primers see Table S14) and used for transfection of NIH3T3 cells. Briefly, cells were seeded in 10 cm dishes and cultivated 24 h to reach 70-90% confluence. For transfection, 10 µg purified plasmid DNA, 20 µl TurboFect, and 1 ml DMEM were combined and vortexed immediately. Then, the mixture was incubated 20 min at RT, and added dropwise to the cells. Cells were harvested by scraping in ice cold 1x PBS, pelleted by centrifugation at 500 g, 4° C for 5 min and flash frozen using liquid N_2_.

### SDS–PAGE and immunoblotting

Tissues/cells/neurospheres/oligospheres were resuspended in 4% SDS, 0.1 M HEPES pH 7.4, incubated at 95° C for 10 min, and sonicated for 1 min at an amplitude of 60% (UP50H, Hielscher, Teltow, Germany). For neurosphere and oligosphere samples, incubation at 95° C was repeated after sonication. Lysates were centrifuged at 20,000 g, RT for 30 min, the clear supernatant transferred to a new tube, and the protein concentration determined using the DC protein assay (BioRad, Hercules, CA). Equal amounts of protein were combined with 4x sample loading buffer (4% β-mercaptoethanol, 8% SDS, 40% glycerol, 4% bromophenol blue, 240 mM Tris-HCl pH 6.8) to a final concentration of 1x, incubated for 10 min at 95° C, and proteins were separated using SDS–PAGE. Proteins were blotted to 0.45 µm nitrocellulose membranes and blocked using either 5% non-fat milk or 3% BSA in Tris-buffered saline with 0.05% Tween 20 (TBS-T). The following primary antibodies were incubated with the blots overnight at 4 °C in blocking solution: anti-myc-tag (1:5,000, rabbit pAb, ab9106, Abcam, UK), anti-Dclk1 (1:1,000, rabbit pAb, ab31704, Abcam, UK), anti-Padi2 (1:1,000, rabbit pAb, 12110-1-AP, Proteintech Group, Rosemont, IL), anti-Pura (1:1,000, rabbit pAb, ab79936, Abcam, UK), anti-FLAG-tag (1:5,000, mouse mAb, F1804, Sigma-Aldrich, St. Louis, MO), anti-alpha-tubulin (1:2,000, rabbit pAb, 600-401-880, Rockland Immunochemicals, Gilbertsville, PA), and anti-beta-actin (1:5,000, mouse mAb, A5316, Sigma-Aldrich). Blots were washed thrice with TBS-T and incubated with one of the following secondary antibodies for 1 h at RT: HRP-goat anti-rabbit (1: 5,000, goat pAb, 111-035-003, Dianova, Germany) or HRP-goat anti-mouse (1:5,000, goat pAb, 115-035-044, Dianova, Germany). For detection of protein biotinylation, membranes were incubated with HRP-Streptavidin (1: 10,000, 21126, Thermo Fischer Scientific) in 3% BSA, 1x PBS-T and washed 6x with 1x PBS. Specific binding was detected by enhanced chemoluminescence using Clarity Western ECL Substrate (BioRad, Hercules, CA) and visualized with a FUSION SOLO 4M system (Vilber Lourmat, Eberhardzell, Germany).

### MS sample preparation, TMT labeling, and peptide fractionation

Neurospheres were resuspended in ice cold lysis buffer (0.5% NP-40, 50 mM HEPES, 10% Glycerol, 150 mM KCl, 1 mM MgCl_2_, and 1x cOmplete protease inhibitor cocktail (Roche Diagnostics, Mannheim, Germany)) at a pellet to buffer ratio of 1:10 (v/v) and lyzed by sonication with 3x30 sec cycles at an amplitude of 80% on ice. After centrifugation at 20,000 g, 4° C for 30 min, the supernatant was transferred to a new tube, the protein concentration determined using the DC protein assay, and 130 µg of protein were precipitated by Cl/MeOH followed by in solution digested as described elsewhere.^22^ Briefly, protein pellets were resuspended in 0.1 M TEAB/0.1% RapiGest (Waters, Milford, MA), reduced with 5 mM DTT and alkylated with 20 mM acrylamide.^23^ The reaction was quenched by addition of 5 mM DTT and 1.5 µg of trypsin (Promega, Madison, WI) was added to the sample. Digestion was carried out overnight at 37° C followed by TMT 6plex labeling the following day. TMT labels (Thermo Fisher Scientific) were dissolved in 41 µl ACN, added to the respective samples, mixed by vortexing, and incubated for 90 min at RT. The reaction was stopped by addition of 8 µl 5% hydroxylamine in 0.1 M TEAB pH 8.5 and incubation for 15 min at RT. After combination of samples, the amount of acetonitrile was reduced to <5%, TFA added to a final concentration of 0.1%, and the combined sample desalted using 10cc Oasis HLB cartridges (Waters). The eluate fraction was dried using a vacuum centrifuge and resuspended in 3.6 mL ampholyte solution (pI 3-10, GE Healthcare, Chicago, IL). Offgel fractionation was performed using a 3100 OFFGEL fractionator (Agilent, Santa Clara, CA) in the 24-fraction setup using pI 3-10 IPG strips (GE Healthcare) for 50 kVh. Individual fractions were desalted using Stage Tips^24^ and dried using a vacuum centrifuge.

### Dimethyl labeling, SCX fractionation, and phosphopeptide enrichment

Neurosphere lysis, digestion and strong cation exchange (SCX) fractionation were performed as described elsewhere.^25^ Briefly, neurosphere pellets were resuspended in 10 volumes of ice-cold lysis buffer (8 M urea, 75 mM NaCl, 50 mM HEPES pH 8.2, 1 mM sodium fluoride, 1 mM β-glycerophosphate, 1 mM sodium orthovanadate, 10 mM sodium pyrophosphate, 1 mM PMSF, and 1x cOmplete protease inhibitor) and immediately mixed by pipetting followed by sonication for 2× 30s at 80% amplitude on ice. Lysates were centrifuged for 10 min at 2,500 g, 4 °C and the supernatants were transferred to new tubes. Protein concentration was determined using the DC Protein Assay and 3 mg of starting material were used for each sample. Disulfide bridges were reduced using 5 mM DTT for 25 min at 800 rpm, 56° C in a thermomixer and alkylated using 20 mM acrylamide for 30 min at RT in the dark.^23^ The reaction was quenched by addition of 5 mM DTT for 15 min at RT and samples diluted 1:5 with 25 mM HEPES pH 8.2. CaCl_2_ was supplemented to a final concentration of 1 mM, trypsin was added at an enzyme-to-substrate ratio of 1 to 200 (w/w), and digestion was carried out overnight at 37° C. The next day, 3plex dimethyl labeling was carried out as described elsewhere.^26^ Briefly, for each mg of peptides in the digestion solution, 32 µL 20% formaldehyde and 160 µL 0.6 M sodium cyanoborohydride were added to the samples. The light channel was labeled with CH_2_O/NaBH_3_CN, the medium channel with CD_2_O/NaBH_3_CN, and the heavy channel with ^13^CD_2_O/NaBD_3_CN for 1 h at RT. The reaction was quenched using 23 µL 28% NH_4_OH/mg peptides and acidified with 16 µL FA/mg peptides. Subsequently, the individual channels were combined, desalted using 400 mg Oasis HLB cartridges (Waters), and the eluate fraction dried using a vacuum centrifuge. SCX fractionation was performed using an ÄKTA Purifier system (GE Healthcare) equipped with a 100 mm x 9.4 mm PolySULFOETHYL A column (PolyLC, Columbia, MD) at a flow rate of 2 ml/min at 8° C. The following solvents were used: solvent A: 7 mM KH_2_PO_4_, pH 2.65, 30% ACN; solvent B: 7 mM KH_2_PO_4_, pH 2.65, 30% ACN, 350 mM KCl; solvent C: 50 mM K_2_HPO_4_pH 7.5, 500 mM NaCl. pH values were adjusted by phosphoric acid prior to the addition of organic solvents. Initially, the column was equilibrated with a linear gradient from water to 100% C in 2 min, 100% C for 16 min, and a linear gradient from 100% C to water in 2 min. After priming for 40 min with 100% A, samples (peptides resuspended 500 µl of solvent A) were loaded, followed by a washing step for 4 min at 100% A. Peptides were eluted with a linear gradient from 100% A to 70% A/30% B in 48 min and 0% A/100% B in 2 min followed by 8 min at 100% B, a linear gradient to water in 2 min and finally 8 min of water. In total, 12 fractions (12 ml each) were collected, lyophilized, and desalted using 10 mg Oasis HLB cartridges. The eluate fractions were dried using a vacuum centrifuge and resuspended in 500 µl of 5% TFA, 80% ACN, 1 M glycolic acid.^27^ TiO_2_ beads (Sachtleben, Duisburg, Germany) were added to the sample at a peptide to bead ratio of 1:6 (w/w) and incubated for 15 min at 1,200 rpm, RT. Beads were pelleted by centrifugation for 1 min at 13,000 g, RT and the supernatant was transferred to another tube. The beads were washed sequentially with 1 ml 80% ACN, 1% TFA and 1 ml of 20% ACN, 0.1% TFA followed by drying using a vacuum centrifuge. Phosphopeptides were eluted from the beads by incubation with 200 µl 1% NH_4_OH for 15 min at 1,200 rpm, RT. The supernatant was transferred to a fresh tube, acidified with 10 µl FA, and desalted using 10 mg Oasis HLB cartridges.

### Cell culture, proximity biotinylation, and biotin/phosphopeptide enrichment

NIH/3T3 cells were transfected with the individual BirA*-Dclk1 fusion proteins or an empty vector, after 24 h the medium was changed and biotin was added at a final concentration of 50 µM, followed by an incubation for 24 h. Cells were washed with ice-cold PBS, harvested by scraping, transferred to a microtube, and pelleted by centrifugation for 5 min at 1000 g, 4° C. Cell pellets were resuspended in ice cold lysis buffer (50 mM Tris-HCl pH 7.4, 500 mM NaCl, 2% Triton-X100, 0.4% SDS, 5 mM EDTA, 1 mM DTT, 1 x cOmplete protease inhibitor and 1x PhosSTOP phosphatase inhibitor cocktail (Roche Diagnostics)) in a pellet to buffer ratio of 1:10 (v/v). Samples were sonicated for 40 sec with an amplitude of 60% followed by incubation on a tumbling shaker for 1 h at 12 rpm, 4° C. Subsequently, the lysates were centrifuged for 30 min at 20,000 g, 4° C, the clear supernatants were transferred to new tubes, and the protein concentrations were determined using the DC Protein assay. For each condition, 1 mg of protein were combined for the heavy SILAC labeled sample (control) and the light SILAC labeled sample (transfected with individual constructs) followed by enrichment of biotinylated proteins by Streptavidin Sepharose High Performance affinity resin (GE Healthcare). For all steps, beads were pelleted by centrifugation for 1 min at 3,000 g, RT. Streptavidin beads (150 µl slurry/sample) were washed three times with 150 µl lysis buffer, the sample added to the beads, and the mixture incubated overnight on a tumbling shaker at 12 rpm, 4° C. Beads were washed for 10 min, 800 rpm, at RT with a thermomixer using sequentially 1 ml of the following buffers: twice 2% SDS, 100 mM HEPES, pH 7.6; once 0.1% sodium deoxycholate, 1% Triton-X100, 500 mM NaCl, 1 mM EDTA, 50 mM HEPES, pH 7.5; once 250 mM LiCl, 0.5% NP-40, 0.5% sodium deoxycholate, 1 mM EDTA, 10 mM Tris-HCl, pH 8.1; and twice 50 mM NaCl, 50 mM Tris-HCl pH 7.4. Subsequently, proteins were reduced on the beads using 10 mM DTT, 100 mM HEPES pH 8 for 30 min at 800 rpm, 56° C, and alkylated for 30 min with 25 mM acrylamide at RT. The reaction was quenched by addition of 10 mM DTT and incubation at RT for 10 min. Subsequently, 2.5 µg trypsin/mg protein (amount of input before biotin enrichment) were added and samples adjusted to a final volume of 300 µl with 100 mM HEPES pH 8. Digestion was carried out overnight at 800 rpm and 37 °C in a thermomixer. Subsequently, peptides were extracted two times from the beads with 300 µl 5% ACN, 0.1% FA, and the combined extracts desalted with 10cc Oasis HLB cartridges. Eluate fractions were dried using a vacuum centrifuge and double phosphopeptide enrichment with TiO_2_ was performed as described elsewhere.^28^ Briefly, samples were resuspended in 5% TFA, 80% ACN, 1 M glycolic acid. TiO_2_ beads were added to the sample at a peptide to bead ratio of 1:6 (w/w). The mixture was incubated for 15 min at 1,200 rpm, RT on a thermomixer, the beads pelleted by centrifugation for 1 min at 13,000 g, RT, and the supernatant transferred to another tube (flow through fraction). The beads were washed sequentially with 80% ACN, 1% TFA as well as 20% ACN, 0.1% TFA and dried using a vacuum centrifuge. Phosphopeptides were eluted from the beads with 200 µl 1% NH_4_OH for 15 min at 1,200 rpm, RT, the supernatant was transferred to another tube, acidified with FA, and dried using a vacuum centrifuge. Samples were re-solubilized using 70% ACN, 0.1% TFA, and TiO_2_ beads were added at a peptide to bead ratio of 1:6 (w/w). The beads were washed with 50% ACN, 0.1% TFA, dried using a vacuum centrifuge, and phosphopeptides were eluted from the beads using 1% NH_4_OH, followed by acidification with FA and desalting using Stage tips.^24^ Flow through fractions were dried using a vacuum centrifuge, resuspended in 1% ACN, 0.1% FA, and desalted using 10cc Oasis HLB cartridges.

### LC−MS/MS Analysis

Dried peptide samples were resuspended in 5% FA, 5% ACN or 50 mM citrate^29^ (phosphopeptide enriched samples) and analyzed by UHPLC-MS/MS using either an LTQ Orbitrap Velos in combination with an EASY-nLC 1000 or an Orbitrap Fusion Lumos in combination with an Ultimate 3000 RSLCnano System (all Thermo Fisher Scientific). For both systems, in-house manufactured spray tips were used. Tips were generated from 100 µm ID/360 µm OD fused silica capillaries with a P2000 laser puller (Sutter Instrument, Novato, CA), and packed with 5 µm/1.9 µm ReproSil-Pur 120 C18-AQ particles (Dr. Maisch, Ammerbuch-Entringen, Germany) to a length of 30 cm/50 cm for the Orbitrap Velos/Fusion Lumos, respectively. For Orbitrap Velos analyses, peptides were loaded on the analytical column with solvent A (0.1% FA, 5% DMSO) at a flow rate of 1 µl/min, and separated with 60 min or 90 min linear gradients from 99% solvent A, 1% solvent B (0.1% FA, 5% DMSO, 94.9% ACN) to 65% solvent A 35% solvent B at a flow rate of 400 nl/min. Eluting peptides were ionized in the nanosource of the mass spectrometer in the positive ion mode at a cone voltage of 1.6 kV. Survey scans were acquired in the Orbitrap analyzer from m/z 400 to 1200 with a resolution of 30,000 or 60,000 followed by fragmentation of the 10 most abundant ions by either CID and analysis in the LTQ, or HCD and analysis in the Orbitrap for dimethyl- and TMT-labeled samples, respectively. Only multiply charged ions were selected for fragmentation, the repeat count was set to one and the dynamic exclusion was set to 60 and 90 sec for 60 min/90 min gradients, respectively. For phosphopeptide enriched samples, multi stage activation (MSA) was enabled for neutral-loss of phosphoric acid. For Orbitrap Fusion Lumos analyses, peptides were loaded to the analytical column with 100% solvent A (0.1% FA in water) at a flow rate of 600 nl/min, and peptides separated with 120 min linear gradients from 95% solvent A, 5% solvent B (0.1% FA, 90% ACN) to 65% solvent A, 40% solvent B at a flow rate of 300 nl/min. Eluting peptides were ionized in the positive ion mode at a cone voltage of 2 kV and survey scans acquired in the Orbitrap analyzer from 375 – 1500 m/z at a resolution of 60,000. The most intense ions were fragmented by HCD (charge states between +2 and +7) in the top speed mode (3 sec cycle time) with a dynamic exclusion of 60 sec and fragment ions analyzed in the Orbitrap at a resolution of 30,000. For both systems, prior to loading of phosphopeptide-enriched samples, the analytical column was equilibrated with three injections of 50 mM citrate.

### MS Data analysis

For neurosphere proteome analyses by TMT labeling, data were processed using ProteomeDiscover 2.4.0.305 (Thermo Fisher Scientific) in combination with Mascot 2.6.1 (Matrix Science Ltd, London, UK). After recalibration of precursor masses with the Spectrum Files RC node (MS1/MS2 tolerances: 20 ppm/0.02 Da) using a non-linear regression model, spectra were searched against UniProt *Mus musculus* (55,192 entries, release date 06/2019) in combination with two databases containing common contaminants (cRAP (crapome.org) and MaxQuant contaminants (maxquant.org)) in a reverse-decoy approach. Trypsin/P was selected as enzyme and up to two missed cleavages allowed. Mass tolerances were 10 ppm (MS1) and 20 mmu (MS2). Propionamide (Cys) and TMT6plex (peptide N-Term, Lys) were defined as fixed modifications, and oxidation (Met) as well as acetyl (protein N-term) as variable modifications. Identified peptide spectrum matches (PSMs) were validated by the Percolator node based on q-values and target FDRs of 1%/5% (strict/relaxed). The combined PSMs were aggregated to peptides and proteins according to the principle of strict parsimony and finally filtered at 1% FDR on the peptide and protein level. For peptide/protein quantification, TMT reporter ion signals were extracted at the MS2 level with a tolerance of 20 ppm using the most confident centroid. Subsequently, the following filters were applied for peptides selection: unique and razor; reporter abundance based on: automatic; co-isolation threshold: 30%; average reporter S/N: 10. The resulting protein table was filtered for master proteins, exported, and further analyzed using R version 3.5.3 (R Core Team, R Foundation for Statistical Computing, Vienna, Austria) with the R packages openxlsx (version 4.1.5), amap (version 0.8-18), and limma (version 3.42.2).^30^ After removal of contaminants, protein intensities were log2-transformed and batch effects between replicates reduced by normalization. Each TMTplex was individually normalized with the LOESS method,^31^ followed by a linear regression model for individual protein-based estimation of batch effects and subtraction from log2-intensities. Paired t-tests were performed for day 0 relative to the other time points, considering only proteins with at least three valid values per group. P-values were adjusted for multiple testing using the Benjamini-Hochberg (BH) procedure^32^ and values < 0.05 were considered to be significant. Fold changes were calculated as mean difference between sample pairs. For proteins with a significant result in at least one comparison (adjusted p-value < 0.05), mean log2-intensities were calculated across all replicates of the same day and z-scored. These values were used for cluster analysis (hierarchical clustering with complete linkage and Spearman correlation distance). For each cluster, gene ontology (GO) analyses were performed for biological process, cellular component, and molecular function using topGO (version 2.38.1) with all proteins in the data set as the background. For each GO term, fold-enrichment values were calculated as the frequency ratio of the term in the cluster and the background. Graphics were created using the R packages ggplot2 (version 3.3.1),^33^ gplots (version 3.1.1) and scales (version 1.1.1). To visualize regulation of proteins related to the categories “transcription” and “kinase”, UniProt keywords were added via Perseus and line plots generated for individual keywords in GraphPad Prism.

Raw files of dimethyl labeling neurosphere phosphoproteome analyses were processed with MaxQuant^34^ version 1.6.14.0 in combination with UniProt *Mus musculus* (63,666 entries including isoforms, release date 04/2020) and MaxQuant’s internal contaminant database with the following settings: precursor ion tolerance (first/main search): 20/4.5 ppm, MS/MS tolerance (ITMS): 0.5 Da, peptide/protein FDR: 0.01, site FDR: 0.01, minimal peptide length: 7, minimal score for modified peptides: 40. The enzyme specificity was set to Trypsin/P (specific or semi-specific) with two missed cleavage sites. Acetylation (protein N-term), carbamylation (N-term), oxidation (Met) and phoshospho (Ser/Thr/Tyr) were defined as variable modifications, as well as propionamide (Cys) as fixed modification. For quantification, 3plex dimethyl-labeling was defined and both the re-quantify and the match between runs option were enabled with default settings. MaxQuant txt output files (“ProteinGroups” and “Phospho(STY)Sites”) were used for further bioinformatics analysis with Perseus^35^ 1.6.15.0 and R 4.1.2 (+ tidyverse package)^36, 37^. The protein groups table was filtered for contaminant, identified by-site, and reverse hits. Afterwards, normalized abundance ratios were log2-transformed and proteins without quantitative information removed from the dataset. Contaminants and reverse hits were removed from the phosphosites table, followed by filtering for class I sites (localization probability >0.75). Abundance information derived from singly, doubly and multiply phosphorylated peptides was separated (multiplicity splitting), log2-transformed, and sites without quantification removed. Finally, regulated phosphosites were defined for individual comparisons (day 7/ day 0, day 14/ day 0, day 14/ day 7) based on a log2 fold change cut-off of +/-1 in both replicates. To gain further insights into regulated phosphosites, annotation enrichment analyses of UniProt keywords were conducted using the Fisher exact test (Benjamini-Hochberg, FDR: 0.02, relative enrichment: protein) with the whole dataset as background. In addition, sequence pattern analyses were performed using IceLogo^38^ in combination with the precompiled Swiss-Prot *Mus musculus* reference set as background. Line plots for proteins with the UniProt keywords “transcription” and “kinase” were generated in GraphPad Prism.

For SILAC data isoform protein-protein interaction analyses of BioID Dclk1 fusion proteins, raw files were analyzed with MaxQuant against the same database as described above including sequences of myc-BioID (Plasmid #35700, Addgene, Watertown, MA) and Streptavidin (P22629). The same search settings as used for the dimethyl data were utilized with 2plex SILAC quantification. The resulting proteinGroups and phosphosite txt files were further processed with R 4.2.0 (+ tidyverse package). The protein group table was filtered as follows: proteins with the tag “contaminants”, “identified by-site”, and “reverse” were removed. For the phosphosite table, site entries with the tag “contaminants” and “reverse hits”, as well as entries with a localization probability < 0.75 were removed. Subsequently, signal intensities of light and heavy channels were log2-transformed and the data filtered for proteins/phosphosites with at least 2 valid SILAC light values in any experimental group (BioID construct). Subsequently, missing values were imputed using the lowest value in any sample (“min”-method, imputeLCMD, version 2.1) and light/heavy ratios calculated per replicate. Biotinylated proteins/phosphosites were determined by filtering for all entries with at least two log2 ratios > 1 in any experimental group.

Proteins/phosphosites specific for each BioID-Dclk1 construct were extracted and functionally classified by GO overrepresentation analysis against a global mouse background using clusterProfiler (version 4.4.4).^39^ Enriched terms were filtered for a Benjamini-Hochberg FDR of < 0.2 and a p-value of < 0.05 and redundant terms simplified by sematic similarity (Jaccard similarity, minimal p.adjust > 0.7). For subsequent comparisons of individual BioID-Dclk1 constructs, the SILAC light non-imputed log2 intensities were utilized. The most suitable method for normalization was determined via the NormalyzerDE-package (version 1.14.0),^40^ and samples normalized (protein groups = cyclic LOESS, phosphosites = RLR). Finally, missing SILAC light channel intensities were imputed as mentioned above, and differentially abundant proteins/phosphosites determined by a moderated t-test using linear modeling (LIMMA trend, version 3.52.4). Entries were deemed significantly regulated when they showed an Benjamini-Hochberg-adjusted p-value of < 0.05 and an average log2 fold change > |0.58|. Regulations were visualized in volcano plots and heatmaps using the EnhancedVolcano (version 1.14.0) and the pheatmap-package (version 1.0.12, Manhattan distance, complete linkage clustering), respectively. For functional analysis, GO-annotation overrepresentation analyses were conducted with clusterProfiler as described above, but with all biotinylated proteins as local background.

## Results

### Differentiating neurospheres consist predominantly of NSCs and OPCs

We generated neurospheres from mouse P0 cortices and differentiated them to oligospheres by addition of B104 cell-conditioned medium for 15 days. To characterize the composition of spheres at individual time points, we performed immunofluorescence (IF) analyses for markers of NSCs, OPCs, and other major cell types of the CNS. We were only able to cultivate dissociated spheres on cover slips up to day 9 of the differentiation process, cells from later time points (12 and 15 days) did not yield viable adherend cultures. Based on the detection of cell-type specific marker proteins, we observed mainly NSCs and OPCs, as well as small numbers of astrocytes in all preparations, but did not detect any cells expressing markers for neurons, microglia, or oligodendrocytes (Fig 1A, Fig. S1). For NSCs, OPCs, and astrocytes we quantified the abundance of the individual cell types (Fig 1B), detecting for undifferentiated neurospheres 89% NSCs, which gradually decreased to 75% at day 9. While only 7% of cells were initially expressing the OPC marker protein Ng2, this fraction increased to 58% at day 9, reflecting a continuous increase in OPCs during the transition from neurospheres to oligospheres. This implies, that a certain percentage of cells was positive for both Nes and Ng2, which is in accordance with previous studies showing that OPCs can express both proteins.^41, 42^ Contaminating Gfap-positive astrocytes only contributed a minor fraction (1% to 11%) of cells at all time points.

**Figure 1:**
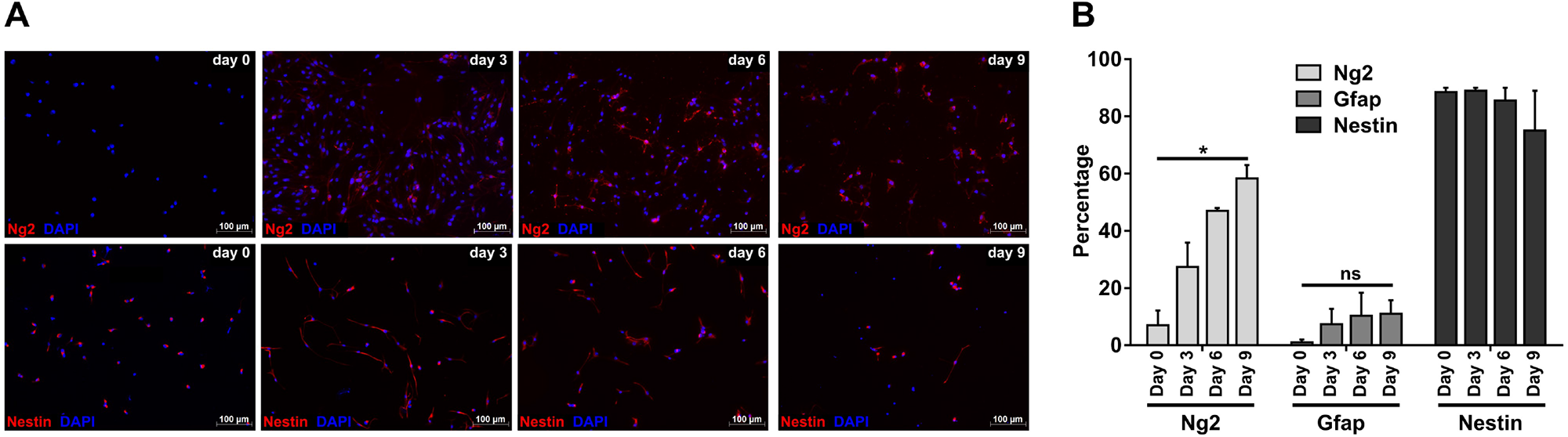
Investigation of the cellular composition of differentiating neurospheres. **A)** Immunofluorescence analysis of dissociated spheres during neurosphere to oligosphere differentiation. Marker proteins shown are Ng2 (OPCs) and Nes (NSCs). **B)** Quantification of cell type abundance during neurosphere to oligosphere differentiation. Marker proteins used: Ng2 (OPCs), Gfap (astrocytes), and Nes (NSCs). For each time point, cell type, and replicate 150 cells each were analyzed. Shown are mean values + SEM, n=3 for Ng2 and Gfap, n=2 for Nes.

### Proteomic analysis of neurosphere to oligosphere differentiation identifies common patterns of protein regulation

For proteomic analyses, we differentiated neurospheres over the course of 15 days to oligospheres in five independent biological replicates. We collected samples every third day, resulting in six time points. After cell lysis, protein extraction, and proteolytic digestion, peptides were TMT-labeled, combined, fractionated by OFFGEL electrophoresis, and analyzed by LS-MS/MS (Fig. 2A). From 373,531 annotated peptide spectrum matches (PSMs), 6,271 protein groups were identified (both at 1% FDR, Fig. S2). Following removal of contaminants and channel-wise normalization, we determined significantly regulated proteins in individual time points relative to day 0 using paired t-tests (Table S1). Based on a combination of an FDR-adjusted p-value (q-value) cut off of 0.05 and a 2-fold change cut-off, we identified consistently increasing numbers of proteins changing in their expression level over the course of differentiation starting at day 9, and reaching a maximum at day 15 with 380 up- and 154 down-regulated proteins, respectively (Fig. S3).

**Figure 2:**
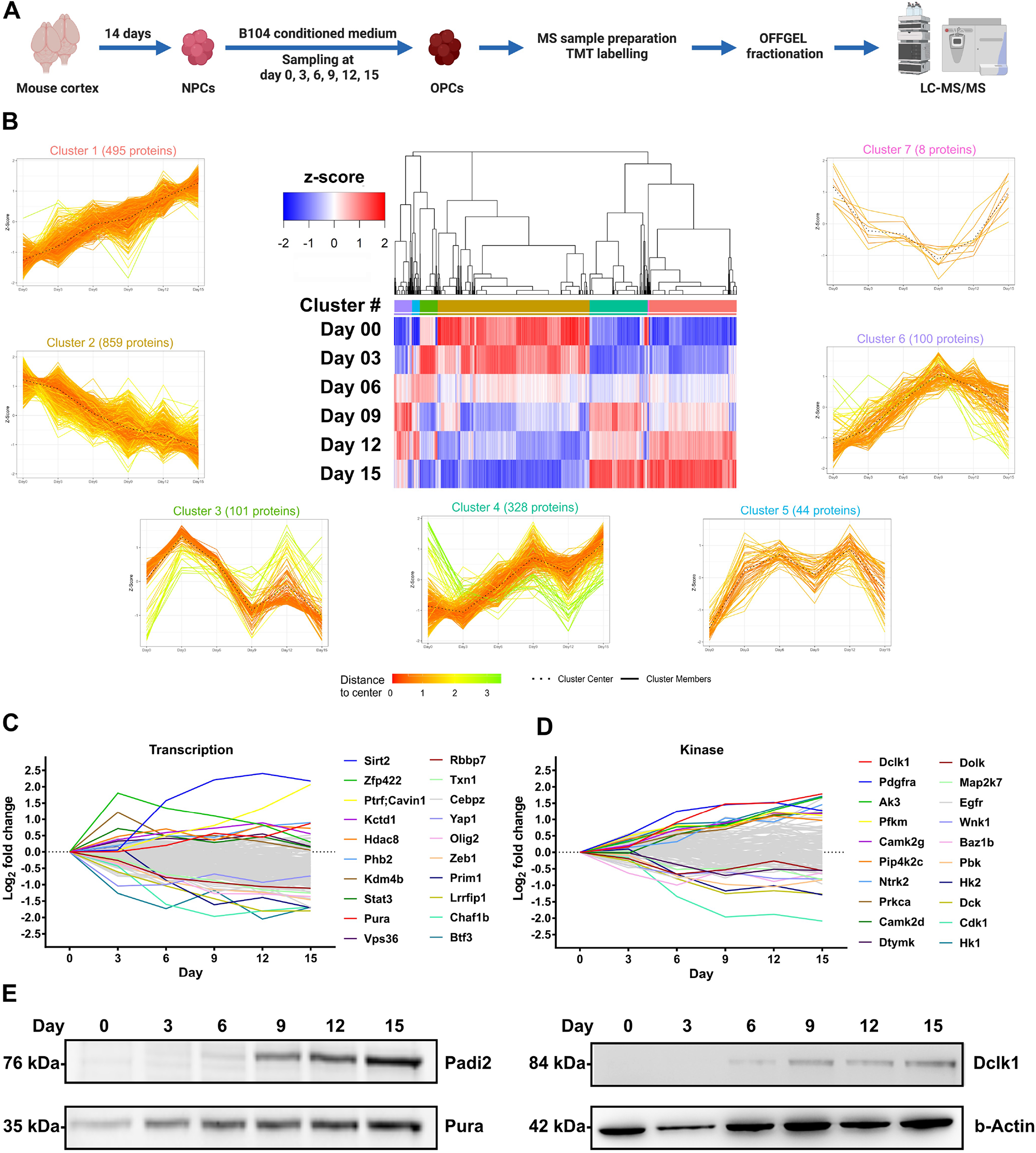
Proteomic analysis of neurosphere differentiation. **A)** Experimental workflow for the proteomic analysis of neurosphere to oligosphere differentiation. **B)** Hierarchical clustering (complete linkage, Spearman correlation distance, based on z-scored mean log2 abundance values, n=7) of the 1935 proteins which were significantly regulated during the differentiation process in at least one time point relative to day 0 (adjusted p-value < 0.05). Based on unsupervised clustering, seven main clusters of similar regulation were observed (patterns of individual proteins are shown as single lines). **C/D)** Average log2 fold-change values of protein subclasses associated with the keywords “kinase” and “transcription”. The 10 most up- and downregulated proteins, respectively, are annotated. **E)** Verification of changes in protein abundance for Padi2, Pura, and Dclk1 by western blot analysis.

To identify groups of proteins with similar regulatory behavior, we performed hierarchical clustering based on time-dependent changes in protein expression during neurosphere to oligosphere differentiation, including all proteins which changed significantly in at least one time point relative to day 0 (q-value < 0.05, Table S1). Subsequently, we assigned individual objects to clusters (agglomerative strategy) based on Spearman correlation distance measures, followed by iterative joining of similar clusters to result in the most compact cluster structure.^43^ This resulted in seven clusters with individual patterns of regulation (Fig. 2B, Table S2), for which we further performed GO analyses (Fig. S4).

Proteins contained in cluster 1 and 4 were upregulated in the majority of time points, implying an OPC-specific expression or a role related to NSC to OPC differentiation. Strikingly, most GO terms enriched in both clusters were related to mitochondria, in particular to the respiratory chain (Fig. S4A/D), and 74% of mitochondrial proteins in the dataset were contained in these two clusters. Cluster 2, which presented the largest group of proteins, contained continuously downregulated proteins. GO analysis identified a strong enrichment in cell cycle regulation and DNA replication, which is reflecting the transition of highly proliferative NSCs to OPCs exhibiting markedly lower division rates.^44^ Furthermore, several complexes related to splicing and RNA binding proteins were included in this cluster, indicating alterations of posttranscriptional regulation of protein expression, a process which was shown previously to be important for OD differentiation.^45^ The other clusters exhibited unique patterns with local maxima/minima. While clusters five, six, and seven did not yield any significant enrichment in the respective GO analyses, cluster three showed a strong overrepresentation of ribosomal proteins and such related to ribosome-related metabolic processes. Proteins contained in this cluster reached maximum intensities at day 3, followed by a continuous decline and a stabilization at later time points. This strong upregulation of ribosomes in the early stages of NSC differentiation could reflect a cellular demand for protein synthesis, enabling proteome remodeling during NSC to OPC proteome transition.

It is well-established, that proteins related to DNA transcription and kinases play an important role in NSC differentiation.^46^ We therefore extracted such annotated with the respective UniProt keywords (226 and 126 proteins, respectively, Figure 2C, D, Table S1). Of the transcription-related proteins, 118 were significantly regulated in at least one time point, and three out of the ten most upregulated hits at day 15 were shown previously to be related to OPCs/ODs: Sirt2, which regulates oligodendroglial cell differentiation;^47^ Pura, a transcriptional regulator of myelin basic protein (Mbp) expression;^48^ and Phb2, which was connected to proliferation regulation of cells of the oligodendroglial lineage.^49^ The top 10 downregulated transcription-related proteins encompassed several members which were implicated in stem cell self-renewal, pluripotency, and (NSC/OD) differentiation, such as Olig2, Yap1, Rbbp7, Lrrfip1, and Zeb1.^50–54^ Out of all kinases contained in the dataset, 42 were significantly regulated in at least one time point. The top ten regulated proteins included Pdgfra, a receptor tyrosine kinase which is known to be crucial for the proliferation of OPCs, as well as Cdk1, Egfr, and Wnk1, which were previously connected to OPC differentiation.^55–58^

Finally, we confirmed the observed expression levels for three selected proteins, namely double cortin like kinase 1 (Dclk1), the transcriptional activator protein Pur-alpha (Pura), and the protein-arginine deiminase type-2 (Padi2) by western blot (Figure 2E), confirming the regulation patterns in our TMT dataset.

### Phosphoproteomic analysis of neurosphere to oligosphere differentiation

As we observed high numbers of regulated kinases, we further investigated changes in protein phosphorylation during neurosphere to oligosphere differentiation (Figure 3A). We collected samples at day 0, 7, and 14, followed by MS sample preparation and 3plex dimethyl labeling.^59^ After pooling of samples, peptides were fractionated using strong cation exchange (SCX) chromatography, phosphopeptides enriched by TiO_2_, and both eluate and flow through fractions analyzed by LC-MS/MS. The resulting dataset contains 4,988 protein groups of which 2,817 were phosphorylated. We identified 9,551 unique phosphopeptides corresponding to 8,608 phosphorylation sites. After removal of contaminants hits, we considered only class I sites (localization probability > 0.75) resulting in 6,398 sites (Table S3, Fig. S5), of which 663 were not reported previously in PhosphoSitePlus.^60^ Subsequently, we identified regulated phosphorylation sites based on individual multiplicities, removing all entries without valid ratios in both biological replicates. This resulted in 300 and 498 phosphosites with a ≥ 2-fold upregulation, and 280 and 512 phosphosites with a ≥ 2-fold downregulation, at day 7 and 14, respectively, relative to day 0 (Figure 3B, Table S3). In contrast to the comparisons with day 0, the number of regulated phosphorylation sites between day 14 and 7 was rather low (57 up- and 38 downregulated), implying that the majority of changes in protein phosphorylation are taking part in the first half of the differentiation process, preceding the changes in protein expression observed in the whole proteome dataset (Figure 2, Fig. S3).

**Figure 3:**
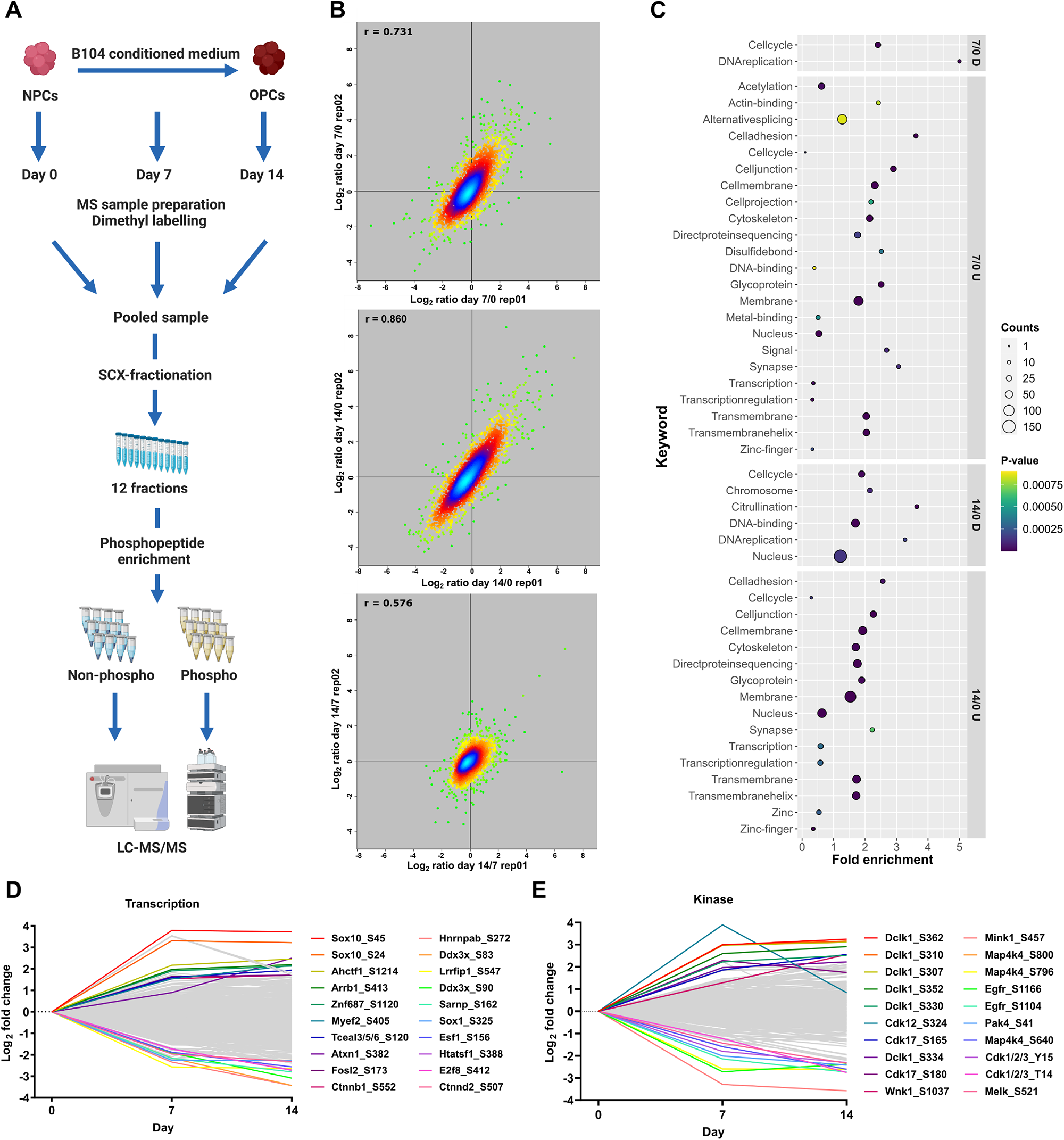
Phosphoproteomic analysis of neurosphere differentiation. **A)** Experimental workflow for the phosphoproteomic analysis of neurosphere to oligosphere differentiation. **B)** Replicate-wise correlation of log2 abundance ratios for individual phosphosites between distinct time points. **C)** UniProt keyword analysis of proteins containing phosphorylation sites significantly regulated between time points (≥ 2-fold up-/downregulated). **D/E)** Average log2 fold-change values of phosphosites contained in proteins associated with the keywords “kinase” and “transcription”. The 10 most up- and downregulated phosphosites, respectively, are shown. D: downregulated, U: upregulated.

We performed UniProt keyword enrichment analyses for proteins containing regulated phosphorylation sites via Fisher’s exact tests (Figure 3C, Table S4), identifying an enrichment of cell cycle regulation proteins in downregulated phosphorylation sites, concurrent with their depletion in upregulated sites. In accordance with this finding, kinase motif analysis of individual phosphorylation sites by Fisher Exact Annotation enrichment for Linear motifs (Table S5) revealed cycline dependent kinase (CDK) motifs as top regulated categories of the downregulated phosphorylation sites of both day 7 and day 14 relative to day 0, while no such pattern was observed for upregulated sites. Furthermore, out of the 33 phosphorylation sites identified for CDKs, 14 were regulated by ≥ 2-fold in at least one comparison, which is in accordance to Cdk1’s downregulation on the protein level and a switch from proliferation to differentiation during the transition from NSCs to OPCs. Next, we analyzed the amino acid sequences adjacent to regulated phosphosites (Fig. S6). While for both the up- and downregulated sites, mainly SP sites with a basic residue in the -2/-3 position were enriched, we furthermore observed an overrepresentation of A in position -1 when comparing upregulated proteins between day 14 and day 7.

Analogous to the TMT dataset, we further manually investigated proteins assigned with the keyword transcription and kinase, for which we identified 707 and 295 class I phosphorylation sites, respectively (Figures 3D, E). The top ten upregulated phosphorylation sites of transcription-related proteins contained four sites located at proteins known to be related to cells of the oligodendroglial lineage: Sox10, which is essential for transcriptional control of oligodendrocyte development,^61^ Myef2, which was shown to control OPC differentiation,^62^ and Ctnnb1, which was found to play a role in Wnt signaling-dependent regulation of oligodendrocyte development.^63^ The top 10 downregulated sites related to this keyword included such located in Hnrnpab, which is involved in neural stem cell differentiation and myelination in ODs,^64^ and Sox1, whose downregulation was shown to lead to enhanced OPC production. ^65^

The top ten regulated phosphorylation sites in kinases originated from only four proteins, Dclk1 (6 phosphorylation sites), Cdk12/Cdk17 (3 phosphorylation sites), and Wnk1. While Dclk1 upregulation was in agreement with protein levels, Wnk1, which was the only protein among the top ten regulated ones shown previously to be related to OPC differentiation,^58^ presented with an inverse behavior (upregulation of protein phosphorylation and downregulation of protein levels). With respect to the downregulated sites, we found such on the OPC-related proteins Egfr and Cdk1 to behave similar to the whole protein levels. Additionally, we detected one site each in Melk and Pak4, which were both found to play a role in NSC proliferation/differentiation.^66, 67^

### Dclk1 isoform abundance correlates with differentiation

Both, in the proteomic and the phosphoproteomic dataset, we found peptides originating from Dclk1 (double cortin like kinase 1) to be highly upregulated during neurosphere to oligosphere differentiation. Dclk1 consists of two N-terminal doublecortin (Dcx) domains, which are connected by an SP-rich region to a C-terminal kinase domain with homology to Ca^2+^/calmodulin-dependent kinases (Camk), but no regulation through calmodulin.^68^ Four main variants were reported for Dclk1 to date, with Dclk1 long and short being most abundant (Fig. S7A).^69^ While Dclk1 long locates mainly to microtubules via its Dcx domains, its C-terminal kinase domain (Dclk1 short) was shown to locate mainly to the nucleus.^70^ Expression of Dclk1 was investigated e.g. in neurons and pancreatic cancer stem cells,^71, 72^ while, to our knowledge, expression in cells of the oligodendroglial lineage has not been reported previously.

Initially, we confirmed changes in Dclk1 expression levels during neurosphere to oligosphere differentiation via western blot (Fig. 4A). While the intensity profile of Dclk1 long correlated with the proteomics data (Fig. 4B), we observed an inversely regulated abundance profile for Dclk1 short. Densitometric quantification revealed a gradual increase of Dclk1 long during the differentiation from NSCs to OPCs, while the abundance of Dclk1 short decreased at the same time (Fig. 4C). This indicates a change in distribution between individual Dclk1 isoforms during the differentiation process. Accordingly, manual investigation of individual peptides revealed that only peptides located in the Dcx domain (which is exclusive for Dclk1 long), were found to continuously increase during the differentiation process (Fig. S7B).

**Figure 4:**
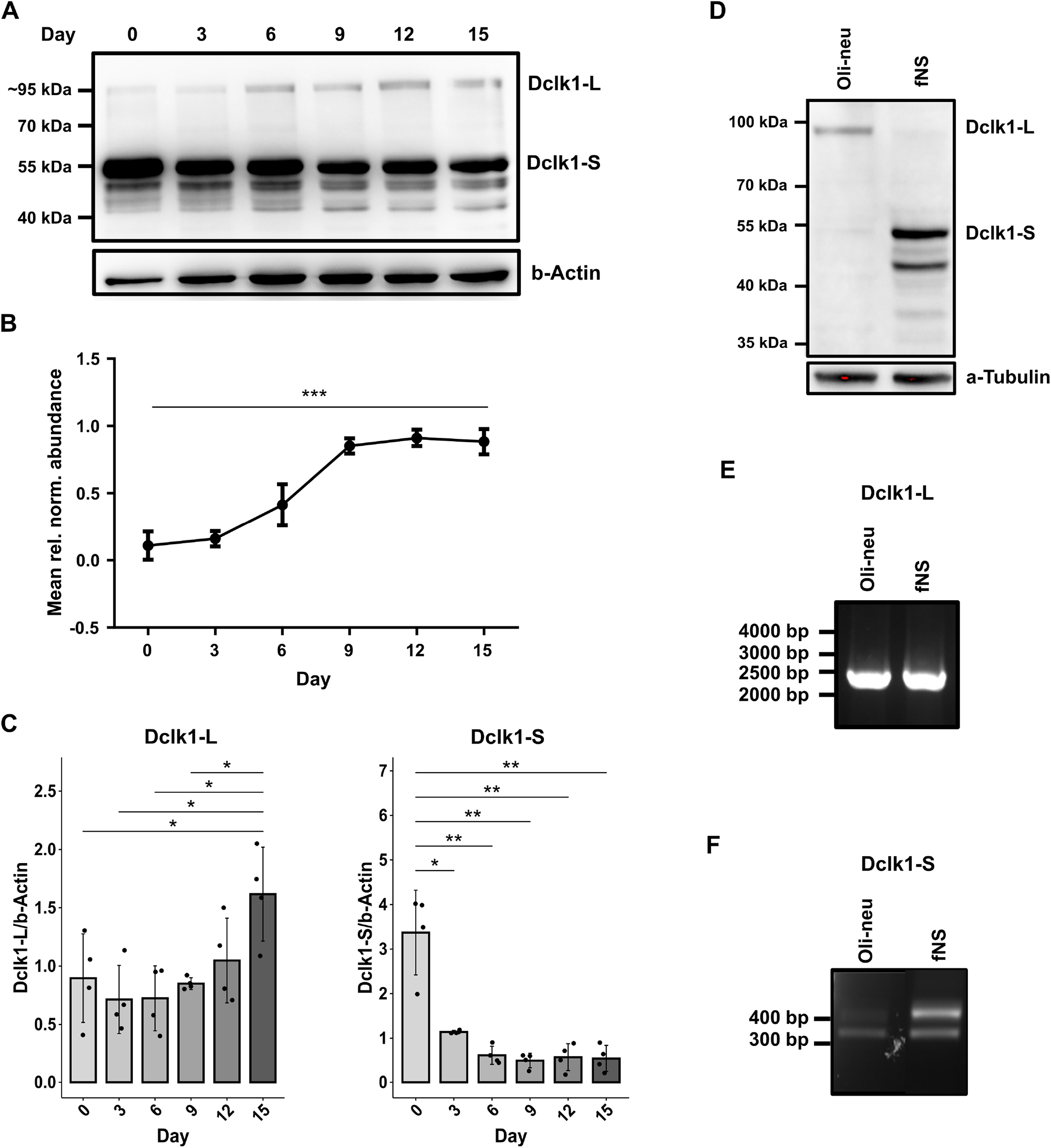
Differential abundance of Dclk1 isoforms in NSCs and OPCs. **A)** Western blot of analysis of Dclk1-expression during neurosphere to oligosphere differentiation. Two distinct isoforms are observed at ∼95 kDa and ∼55 kDa. **B)** MS-based normalized Dclk1 abundance during neurosphere to oligosphere differentiation. Mean scaled peptide intensities (n = 7) of 5 unique Dclk1 peptides ± SEM. *** p < 0.001. **C)** Densitometric quantification of western blot analyses of the abundance of individual Dclk1-isoforms during neurosphere to oligosphere differentiation. Different exposure times were used for individual isoforms (see supporting file 1 for uncropped blots). Individual values were normalized to β-actin. Shown are mean values (n = 4) ± SD; statistical significance was determined using Welch’s t-test; * p < 0.05, ** p < 0.01. **D)** Western blot analysis of Dclk1-isoforms in Oli-neu and fNS cells. **E/F)** RT-PCR mRNA expression analysis of Dclk1-isoforms in Oli-neu and fNS cells. Dclk1-L: Dclk1-long, Dclk1-S: Dclk1-short.

In neurosphere to oligosphere differentiation, NSCs and OPCs present the start/end point of the process. As surrogate for pure populations of these cell types, we utilized fNS and Oli-neu cells,^19, 20^ and investigated the presence of Dclk1 isoforms. Dclk1 short was detected almost exclusively in fNS cells, while the same was true for Dclk1 long in Oli-neu cells (Fig. 4D). This implies that the individual isoforms are specific for NSCs and OPCs, respectively, and that the gradual transition observed in our dataset is related to changing amounts of the individual cell types during the differentiation process. This observation may reflect the cellular differentiation status in general, as it was shown that Dclk1 short is a marker for pluripotency in pancreatic cancer cells,^72^ or may be related to commitment towards the oligodendroglial lineage. We therefore investigated various cell types as well as cortical tissue (the starting material for neurosphere generation) in order to assess if the presence of Dclk1 long/short varies between cell types (Fig. S7C, D). For mouse cortex, B104 neuroblastoma cells, and HEK293 cells, we predominantly observed Dclk1 long, while C6 glioblastoma, B35 neuroblastoma, and NIH/3T3 cells expressed both isoforms. This indicates that presence and abundance of Dclk1 varies across different types of cells, irrespective of their relation to NSCs or the oligodendroglial lineage.

Individual Dclk1 isoforms can be formed by two different mechanisms: alternative splicing, which can result in four different major versions of Dclk1 (Fig. S7A), and proteolytic cleavage, which was shown to generate Dclk1 short from a Dclk1 long precursor.^70, 72^ In order to delineate which mechanism was responsible for the presence of the individual isoforms in fNS and Oli-neu cells, we investigated the mRNAs corresponding to individual splice variants using RT-PCR.^73^ In both fNS and Oli-neu cells, we detected equal mRNA levels of Dclk1 long (Fig. 4E), while Dclk1 short was more abundant in fNS cells (Fig. 4F). While the latter is in accordance with the pattern observed by western blot (Fig. 4D), the absence of the Dclk1 long protein in fNS cells contradicts its mRNA levels. This implies that Dclk1 long is expressed in fNS cells, but is proteolytically cleaved to result in Dclk1 short, which is therefore generated through two mechanisms.

### Phosphorylation of Dclk1 in the SP-rich domain affects its proteolytic processing

Dclk1 was shown to be proteolytically processed by calpain at two different cleavage sites located adjacent to the SP-rich domain.^70^ In our phosphoproteomics dataset, we identified 16 class I phosphorylation sites on Dclk1 (Table. S3). Nine of them were located in the highly conserved region (between mouse, rat, and human) of the SP-rich domain (AA 288-342),^74^ and up to 8-fold upregulated during neurosphere to oligosphere differentiation (Fig. 5A). The concurrent increase of Dclk1 long and phosphorylation of the SP-rich domain could either be due to inactivation of the cleaving protease followed by phosphorylation; or by regulation of proteolytic cleavage by phosphorylation.^75^

**Figure 5:**
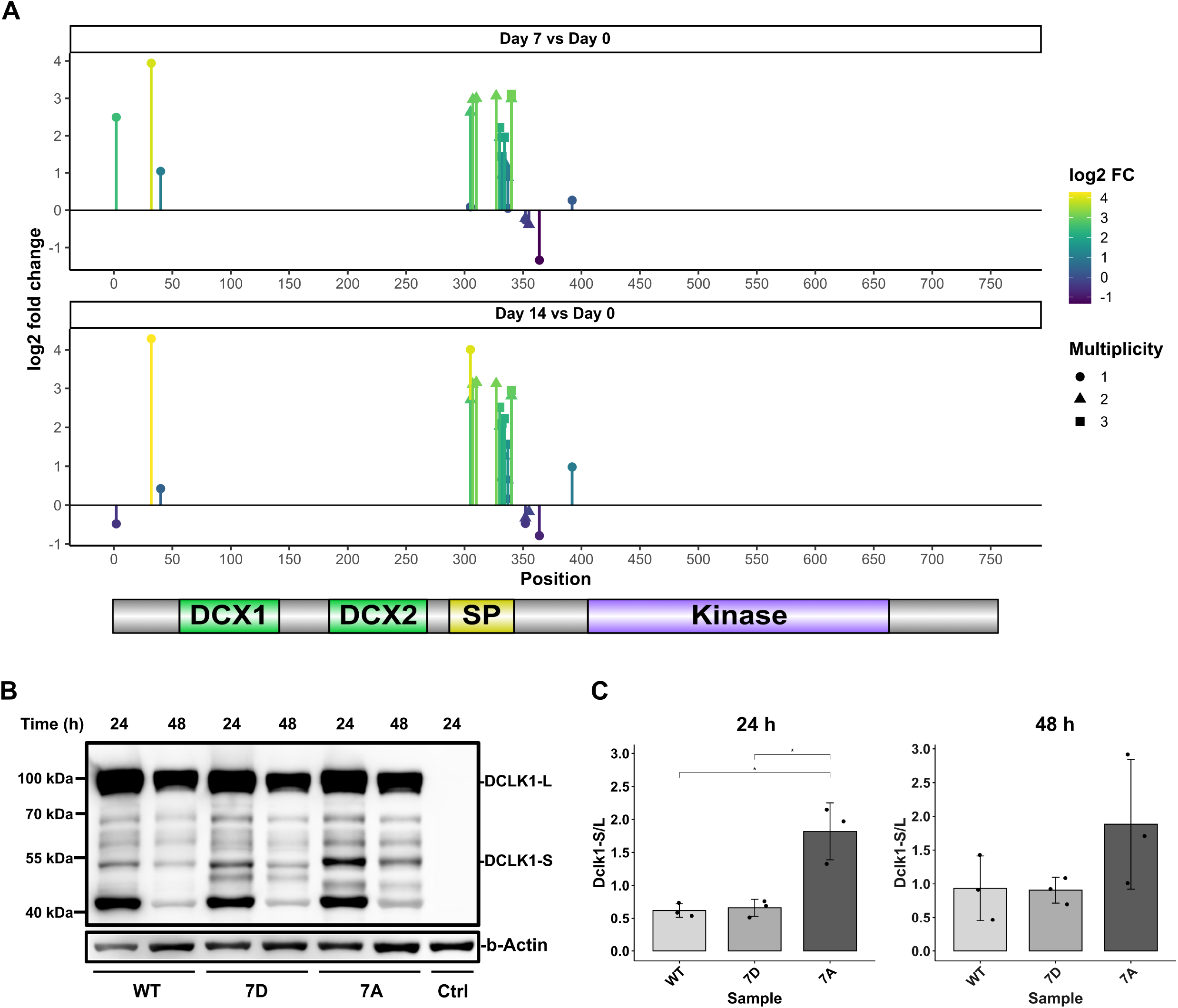
Phosphorylation in the SP-rich region regulates Dclk1 cleavage. **A)** Distribution of quantified Dclk1 class I phosphorylation sites and regulation between individual time points of neurosphere to oligosphere differentiation. Shown are mean log2 fold change values (n = 2). **B)** Western blot analysis of doxycycline induced pulse-chase expression for WT-, 7D- and 7A-Dclk1 after 24 h and 48 h in NIH/3T3 pTR3G cells. Individual constructs were detected via a C-terminal Flag-tag. ctrl.: mock-transfected cells **C)** Phosphomimetic Dclk1 versions (7D) show reduced proteolytic cleavage of Dclk1 in the SP-rich region. Shown are mean values (n = 3) + SD; statistical significance was determined using Welch’s t-test; * p < 0.05.

To differentiate between these possibilities, and to investigate a possible effect of phosphorylation on proteolytic Dclk1 cleavage, we generated phosphomimetic (7S to 7D) and – resistant (7S to 7A) versions of Dclk1 long for the residues we found to be differentially phosphorylated in the SP rich region (S305, S307, S330, S332, S334, T336, and S337). To enable discrimination of endogenous Dclk1, we fused myc- and FLAG-tags to the constructs’ N- and C-termini, respectively. Constructs were expressed in NIH/3T3-pTRE3G cells, since these cells contained both Dclk1 long and short (Fig. S7D), implying that they are able to proteolytically process Dclk1. Following pulsed Dclk1 expression through limited application of doxycycline, we detected Dclk1 fragments of the 7A and the 7D Dclk1 versions after chase times of 24h and 48h by western blotting (Fig. 5B). While the phosphomimetic mutant (7D) resulted in stabilization of Dclk1 long, the phosphoresistant version (7A) showed increased proteolytic cleavage and consequentially release of Dclk1 short (Fig. 5C). Furthermore, we identified varying molecular weights for Dclk1 short for the 7A and 7D version, possibly indicating differences in calpain cleavage^70^ based on the phosphorylation status.

### Identification of potential Dclk1 isoform-specific interaction partners and substrates by proximity biotinylation

The regulation of Dclk1 proteolytic processing during neurosphere to oligosphere differentiation implies distinct functions of the individual isoforms. This could be related to phosphorylation of substrates by the kinase domain present in both the long or the short isoform, or to interaction with different domains (Fig. S7A). As it was shown that Dclk1 long primarily locates to microtubules, while Dclk1 short is present in both the cytoplasm and the nucleus,^70^ individual isoforms are likely to form spatially restricted interaction networks with a possible function in cellular fate.

To investigate the interactome of individual Dclk1 isoforms, we generated four BioID^76^ constructs fusing the biotin ligase BirA* to the N-terminus of Dclk1 long (DSK), its kinase domain (K), the SP rich + kinase domain (SK), as well as the Dcx + SP rich domain (DS, Fig. 6A). We expressed the individual fusion proteins in NIH/3T3 cells, induced biotinylation, and confirmed construct expression as well as interaction partner biotinylation by western blotting (Fig. 6B, C). For mass spectrometric analyses, we included heavy SILAC labeled control cells to allow for discrimination between biotinylated and background proteins. Subsequently, we performed streptavidin affinity enrichment, on bead digestion, and phosphopeptide enrichment, followed by LC-MS/MS analysis of individual fractions (Fig. 6D).

**Figure 6:**
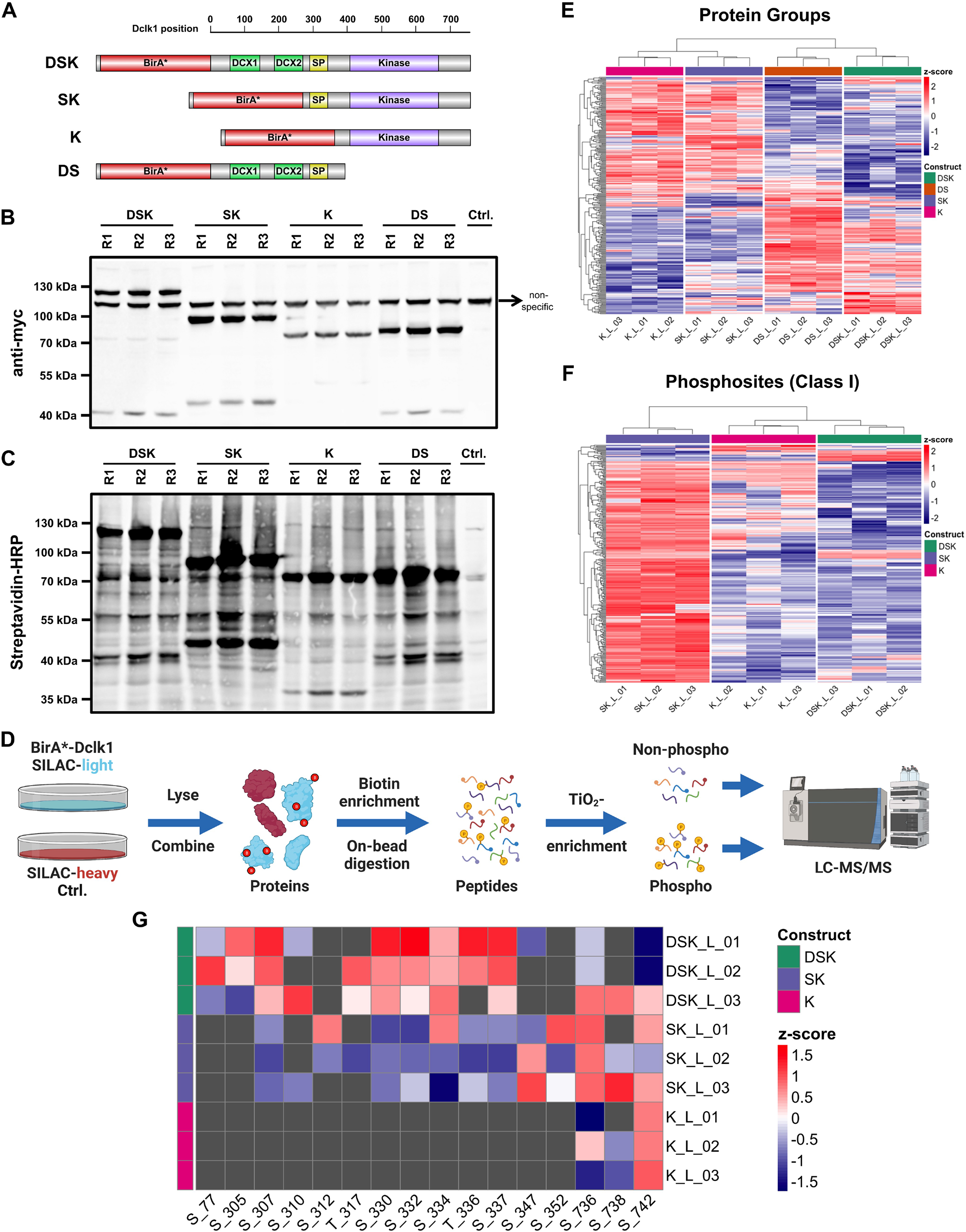
Investigation of Dclk1 isoform-specific interactomes by proximity biotinylation. **A)** Fusion proteins of BirA* and different Dclk1-domains for proximity biotinylation experiments. **B)** Investigation of fusion protein expression in NIH/3T3 cells via the constructs’ N-terminal myc-tag. **C)** Investigation of proximity biotinylation for individual fusion proteins in NIH/3T3 cells. **D)** Workflow for (phospho-) proteomic analysis of Dclk1-BirA* fusion protein interactomes. **E)** Intensity distribution of protein groups identified for the four individual Dclk1 constructs to be significantly regulated relative to another construct. **F)** Intensity distribution of phosphosites identified for the four individual DCLK1 constructs to be significantly regulated relative to another construct. **G)** Z-scored log2 intensities of Dclk1 phosphorylation sites in individual constructs. D: Dcx domain; K: kinase domain; S: SP-rich domain.

### Microtubule-bound and cytosolic/nuclear Dclk1 isoforms form unique interaction networks

For the interactome dataset, we identified 3,424 proteins, of which 1,779 were enriched relative to the heavy SILAC labeled control (Table S6, Fig. S8). For those proteins, we performed principle component analysis (PCA) and unsupervised hierarchical clustering/heatmap analyses, which indicated a clear difference in the interactome of the individual constructs (Fig. S8C, D). We observed a clear separation of constructs containing the Dcx domain (DS and DSK) from such lacking it (K and SK), demonstrating distinct interactomes. To follow up on this, we performed GO enrichment analysis for individual constructs, identifying as top enriched categories microtubule-related processes, splicing/mRNA processing, the proteasome, and myelin sheath (Fig. S9).

We then matched individual datasets, identifying 282 proteins to be differentially abundant in at least one pairwise comparison (Table S8, Fig. S10). Heatmap analyses of these proteins revealed that such interacting with constructs of a similar subcellular localization also showed similar patterns (Fig. 6E). Interestingly, presence of the SP rich domain had only minor effects on the interactome of the kinase domain, which was underlined by the direct comparison of these two datasets (Fig. S10A). For the comparison of microtubule-localized constructs (DSK vs. DS), we found slightly higher numbers of interaction partners for DS, which is lacking the kinase domain (Fig. S10B). We observed the highest discrepancies when we compared the microtubule-localized to the cytosolic constructs (Fig. 10C-F). GO analyses of proteins with higher signal intensities for microtubule-localized constructs revealed, as expected, mainly categories related to microtubules, as well as mitotic spindles (Fig. S11A), which is in accordance to previous findings.^77, 78^ When considering all individual comparisons, we found 141 proteins which were overrepresented for microtubule-localized constructs (Fig. S11B) and 139 for cytosolic/nuclear fusion proteins (Fig. S11C), of which 67 and 70 were overrepresented in at least 2 comparisons respectively. We matched proteins found to be regulated in all 4 comparisons (21 for microtubule and 17 for cytosolic/nuclear) to such which were altered in our neurosphere to oligosphere differentiation TMT dataset (Fig. 2B, Table S1). Nine of these TMT dataset proteins interacted stronger with the microtubule-localized constructs (DS and DSK, Fig. S11B), and eight with the cytosolic/nuclear fusion proteins (K and SK, Fig. S11C). While the later contained four proteins locating to the nucleus, five of those proteins interacting with the microtubule-localized constructs were related to the cytoskeleton, which is in accordance with the published distribution of individual Dclk1 isoforms. Interestingly, the cytosolic/nuclear group contained Wnk1, which was already identified in the group of the highest regulated kinases, both for the proteomic and phosphoproteomic investigation of neurosphere to oligosphere differentiation (Fig. 1D, 3E), and was shown previously to be involved in the regulation of OPC differentiation.^58^ As we identified both Dclk1 short and Wnk1 to be downregulated during the differentiation process, and we found Wink1 to interact significantly more with Dclk1 short than long, a functional connection could exist.

### Phosphoproteomic analysis of Dclk1 isoform interaction networks reveals potential substrates and subcellular localization-specific phosphorylation patterns of Dclk1 domains

In the phosphoproteomic analyses, we identified 2,391 phosphopeptides from 1,136 proteins, covering 3,677 phosphorylation sites. Of those, 1,952 were classified as class I phosphosites (localization probability >0.75), including 1,048 sites which were enriched in ≥ 2 replicates of the same BioID-construct ≥ 2-fold relative to the control sample (Table S10, Fig. S12). Matching of these phosphosites to such found in the neurosphere to oligosphere differentiation dimethyl dataset (Table S3) revealed an overlap of 618 sites. As we were especially interested in potential substrates of Dclk1, we focused for subsequent analyses on datasets from constructs containing the Dclk1 kinase domain (DSK, SK, and K).

Global comparison of all phosphorylation sites found to be significantly enriched relative to the heavy SILAC labeled control samples by PCA and unsupervised hierarchical clustering/heatmap analyses revealed a clear separation of the individual constructs’ phosphoproteomes (Fig. S12C, D). GO analyses of individual datasets identified similar categories as such performed on the protein level, with the major discrepancy that no subunits of the proteasome were identified, but proteins related to histone modification, which were strongly overrepresented for K and SK (Fig. S13). Interestingly, this contained also categories related to mRNA processing and splicing, which is in accordance with a recent study investigating putative Dclk1 substrates.^79^ Subsequently, we directly compared phosphosites for the different constructs. BirA*-SK yielded the largest number of overrepresented phosphorylation sites relative to the other fusion proteins (Fig. 6F, Fig. S12E). In these binary comparisons for all datasets, intensities of phosphorylated peptides were markedly increased for SK relative to K (Fig. S14A), while these constructs showed very similar values for their interactome (Fig. 6E), implying a potential regulatory function of the SP-rich region with respect to protein phosphorylation. This included, for example, S88 in Lrrfip1, a protein which we also identified as one of the top regulated proteins involved in transcriptional regulation in the TMT dataset, and which was connected previously to NSC differentiation (Fig. 3D).^50^ Also, for the comparison of SK and DSK we observed a similar trend, while comparison of the latter with K only showed a small number of differentially abundant phosphorylation sites (Fig. S14B, C).

Finally, we matched phosphorylation sites which were regulated at day 7 or day 14 relative to day 0 in the dimethyl dataset to such identified in the Dclk1 BioID experiments, identifying 131 shared sites (Fig. S14D). We then individually matched experiments based on the expression of Dclk1 short/long during neurosphere to oligosphere differentiation. Consequentially, we compared SK/K specific sites with such downregulated during neurosphere to oligosphere differentiation and DSK sites with such found to be upregulated (Table S3, S12). Of the microtubule-bound/dimethyl-upregulated sites (Fig. S14E), five were related to Dclk1’s SP rich region, confirming the correlation between phosphorylation in this part of the protein and the increase in Dclk1 long abundance during NSC to OPC differentiation. Additionally, S517 on microtubule-associated protein 4 (Map4) and S454 on Map3k4 matched between both datasets. Interestingly, inactivation of Map3k4 was connected to regulation epithelial-to-mesenchymal transition and cellular stemness,^80^ providing a possible connection to the differential abundance of Dclk1 short/long in NSCs/OPCs, respectively. For cytosolic-nuclear/dimethyl-downregulated sites (Fig. S14F), we identified seven phosphorylation sites in different proteins (S176 on Acin1, S1870 on Akap13, S133 on Eef1d, S1969 on Jmjd1c, S88 on Lrrfip1, S216 on Mllt4, and S2624 on Srrm2). In accordance with a recently published study identifying mRNA-related processing-related proteins as putative substrates of Dclk1,^79^ five of these proteins were shown to be related to such functions, including splicing (Acin1 and Srrm2),^81, 82^ translation (Eef1d),^83^ and transcriptional regulation (Lrrfip1 and Jmjdc1).^84, 85^ Furthermore, Mllt4/AFDN was shown to be involved in PDGF signaling^86^ and cell division,^87^ in line with the processes found to be regulated on both the protein and phosphorylation site level in our datasets (Figs 2 and 3).

With respect to Dclk1 itself, we detected a total of 16 phosphorylation sites in the BioID experiments (Fig. 6G). As individual constructs differ in their Dclk1 domains and subcellular localization, we investigated the effect of these factors on the abundance of the protein’s individual phosphorylation sites. In accordance with the dimethyl quantification dataset and its overlap with the BioID experiments (Fig. 5A, S14E), we identified pronounced phosphorylation of the SP-rich region for the microtubule-located full-length protein (DSK), while the cytosolic/nuclear SK samples showed reduced phosphopeptide intensities in this region. This implies that microtubule-bound Dclk1 undergoes a spatially restricted phosphorylation in the SP-rich region, which is in line with impairment of its proteolytic cleavage, and therefore release of the kinase domain. At the same time, both non-microtubule-bound proteins (K and SK), presented with increased phosphorylation levels in the kinase domain. As these sites are located outside of the active center of the kinase domain,^79^ their biological function remains unclear at this point.

Based on the BioID dataset, we further investigated the *in vivo* Dclk1 proteolytic cleavage sites by semi-tryptic searches (Table S13). Focusing on the SP-rich region, we observed two amino acid stretches presenting with a ladder structure (Fig. S15). Such patterns could either originate from proteolytic cleavage without strict amino acid specificity or in-source fragmentation of peptides during ESI. In order to exclude the latter, we matched the retention time profiles of peptides originating from the same region and excluded amino acid stretches directly adjacent to proline. For the first region (Fig. S15A), this resulted in SVNG (residues 313-316) and SQLS (residues 320-323), and for the second region (Fig. S15B) in ISQH (residues 346-349) and LSST (residues 356-359) as possible cleavage sites, respectively. We then matched these results with predicted calpain cleavage sites for Dclk1.^88^ Strikingly, out of five high scoring sites for the SP-rich region (Table S13), two matched with the regions for which we observed semitryptic peptides (S323 and S357, both with a score of 0.96 and p-value < 4.1e-5) while two of the other sites coincided either with a tryptic cleavage site (R342) or were located in a peptide which was below the minimal length for detection (Q384). While none of the predicted cleavage sites was found to be phosphorylated in our datasets, they were located within the phosphorylated region ranging from S305-S352.

## Discussion

In the current study, we utilized the neurosphere assay to investigate NSC to OPC differentiation. To our knowledge, only one other study, which focused on the differentiation of NSCs to neurons, analyzed neurospheres using quantitative proteomics to date, achieving a similar performance with regard to protein identification and quantification.^89^ Initially, we performed IF analyses of dissociated spheres, confirming that the majority of cells expressed markers for NSCs and OPCs with minimal contamination from other glia or neurons, and a gradual transition between cell types during the differentiation process. Both the purity and differentiation kinetics are in line with previous analyses of neurospheres/oligospheres, indicating successful differentiation.^15, 90^ In agreement with a gradual transition between cell types, cluster 2 of our whole proteome TMT-based quantification analyses contained mainly proteins which are related to DNA replication and cell division, whose continuous downregulation is indicative of cell cycle exit. Simultaneously, we observed an upregulation of proteins involved in mitochondrial energy production (cluster 1). This could be related to an increased energy demand during protein/lipid biosynthesis for the pronounced morphological changes observed during OPCs formation. Interestingly, cluster 4, which contains the majority of ribosomal proteins, does not match this pattern of continuous up-/downregulation, and presents only in early time points with high values, followed by a strong decline. This could possibly be due to an initial boost in protein biosynthesis as part of a massive rearrangement of the cellular proteome upon induction of differentiation.

The relatively subtle continuous up-/downregulation of protein levels, with no statistically significant changes at day 3/day 6 relative to day 0, was followed by strong upregulation in the following time points. This is in contrast to our phosphoproteomic analysis, in which phosphorylation site abundance changed dramatically between day 0 and day 7, but remained largely constant for the following week (day 7 relative to day 14). This implies that signal transduction pathways involved in NSC to OPC differentiation were already fully activated at day 7, possibly acting as driving factors of cellular transformation, and retained their activity levels in the following 7 days. Importantly, we identified several proteins which were regulated both on the protein and phosphorylation level. This includes several kinases and transcription-related proteins which were connected previously to the oligodendroglial lineage, as well as such involved in cellular proliferation and differentiation. For example, we identified two strongly upregulated phosphorylation sites on Sox10 (S24 and S45), which is known for its role in oligodendroglial lineage differentiation.^61^ Phosphorylation at these residues, which are highly conserved between mouse and human, was shown to affect stability and transcriptional activity of Sox10,^91^ implying possible roles in the regulation of Sox10 activity during NSC to OPC differentiation. We also detected differential protein abundance/phosphorylation levels of several Cdks, Cdk-interacting proteins, and Cdk inhibitors (Cdk1, 2, 3, 5, 16, and 17 as well as Cdkn1b and Cdkn2aip). Interestingly, it has been shown that regulation of cell cycle and differentiation are interlinked,^92^ implying possible functions for these regulatory events both with respect to NSC proliferation and NSC to OPC differentiation.

One of the most prominently upregulated proteins in both datasets was Dclk1, which belongs to the doublecortin gene family, encompassing at least 11 known paralogues in human and mice.^74^ Dclk1 was shown to occur in various isoforms, which have been related to different cell types and biological functions. For example, Dclk1 long has been shown to regulate neuronal migration and actin structure in a redundant fashion with doublecortin, which shows a high sequence similarity to its N-terminal domain.^71, 93^ Furthermore, it was demonstrated that it is able to control spindle formation during mitosis in the developing brain.^78^ These findings match our BioID datasets, in which we detected proteins related to microtubule depolymerization and spindle formation as highest-enriched GO categories for the comparison of Dclk1 short and long. Furthermore, Dclk1 has been shown to play important roles in various types of cancer. It was, for example, demonstrated that Dclk1 short is the most abundant isoform in colon adenocarcinomas, while normal colon tissues mainly contained Dclk1 long,^72^ and Dclk1 was established as marker for intestinal, renal, and liver cancer.^94–96^ Our observation that Dclk1 short is the dominant isoform in pluripotent highly proliferative NSCs (neurospheres/fNS cells), while the lineage-committed OPCs contained mainly Dclk1 long (oligospheres/Oli-neu cells), is in line with a possible correlation of Dclk1 short and cellular pluripotency/proliferation. At least in our system, the western blot and qPCR analyses indicate that this is based on both transcriptional and proteolytic generation of Dclk1 short.

Our phosphoproteomic data indicate that the proteolysis-dependent Dclk1 isoform transition depends on its phosphorylation status in the SP-rich region, which we could confirm by site-directed mutagenesis experiments. This region, which is frequently also referred to as PEST domain (proline (P)/glutamic acid (E)/serine (S)/threonine(T)-rich domain), was shown previously to facilitate proteolytic cleavage (and degradation) of proteins,^97, 98^ and its phosphorylation has been shown to regulate proteolytic degradation,^99^ further supporting the proposed mechanism. Additionally, it was shown that Dclk1 is cleaved by calpain, facilitating the release of Dclk1 short, and the authors predicted that this proteolytic event takes place in the SP-rich region.^70^ In line with this proposed mechanism, we identified two potential regions of Dclk1 proteolytic processing in the SP-rich region *in vivo*, which matched to bioinformatically predicted calpain cleavage sites.

To identify putative interactors and substrates, we performed proximity biotinylation experiments in combination with (phospho) proteomic analyses for individual Dclk1 isoforms. These analyses revealed individual interactomes for Dclk1 long and short, which, in accordance with the subcellular localization of the individual constructs, were dominated by cytoskeletal and nuclear proteins, respectively. Interestingly, we identified markedly higher numbers of phosphorylation sites for cytosolic/nuclear localized Dclk1 constructs compared to such which are microtubule bound, implying increased kinase activity of Dclk1 short. This could provide a possible explanation for its suggested role in the regulation of cancer stem cell pluripotency,^72^ as its localization to microtubules seems to strongly reduce the phosphorylation of certain substrates in our setup. Along this line, out of the seven proteins which we detected to be phosphorylated in a similar fashion between the whole phosphoproteome and the BioID dataset for Dclk1 short, four were shown previously to be related to cancer. Of those, Lrrfip1/Gcf2, which was also identified as one of the most highly regulated transcription-related candidates on the protein and phosphorylation site level in our large scale datasets, was demonstrated to play a role in colorectal cancer metastasis,^100^ which correlates with the reported overexpression of Dclk1’s short in this type of cancer.^72^

With respect to NSC to OPC differentiation, it is conceivable that the increased kinase activity of Dclk1 short plays a similar role, promoting pluripotency and proliferation, which could play a role in the maintenance of NSC identity.

## Acknowledgements

The authors are grateful to Benjamin Odermatt for valuable discussions as well as Norbert Rösel and Asisa Muchamedin for technical assistance. We would like to acknowledge the following funding agencies: Robert Hardt was funded by the German Research Foundation (DFG) via the research unit FOR2625, Alireza Dheghani was supported by the German Academic Exchange Service (DAAD), and Carmen Schoor by the BONFOR program of the Medical Faculty of the University of Bonn. Karin Schork received funding from CUBiMed.RUB and Martin Eisenacher from the de.NBI project (FKZ 031 A 534A) of the Bundesministerium für Bildung und Forschung (BMBF).

## Author contributions

Robert Hardt: Data Curation, Formal analysis, Software, Visualization, Writing – Review & Editing; Alireza Dehghani: Funding acquisition, Investigation, Validation, Visualization, Writing – Review & Editing; Carmen Schoor: Investigation; Markus Gödderz: Investigation; Nur Cengiz Winter: Investigation; Shiva Ahmadi: Investigation; Ramesh Sharma: Investigation; Karin Schork: Formal analysis, Software, Visualization, Writing – Review & Editing; Martin Eisenacher: Resources, Supervision, Funding acquisition; Volkmar Gieselmann: Conceptualization, Methodology; Dominic Winter: Conceptualization, Project administration, Funding acquisition, Methodology, Writing – Original Draft, Writing – Review & Editing.

## Supplementary Material

### Supplementary Tables and File

**Table S1:** TMT labeling-based quantitative whole proteome analysis of neurosphere to oligosphere differentiation. Filtered protein groups table, differential expression analysis, unsupervised clustering, keyword analyses, Dclk1 peptide analysis.

**Table S2:** TMT labeling-based quantitative whole proteome analysis of neurosphere to oligosphere differentiation. Cluster memberships of individual proteins and gene ontology (GO) analyses of distinct clusters.

**Table S3:** Dimethyl labeling-based quantitative whole (phospho-) proteome analysis of neurosphere to oligosphere differentiation. Class I phosphorylation sites, differential expression analysis, phosphorylation sites with assigned UniProt keywords transcription and kinase, and Dclk1 phosphosite analysis.

**Table S4:** Dimethyl labeling-based quantitative whole (phospho-) proteome analysis of neurosphere to oligosphere differentiation. Unbiased annotation enrichment analysis based on UniProt keywords for regulated class I phosphorylation sites.

**Table S5:** Dimethyl labeling-based quantitative whole (phospho-) proteome analysis of neurosphere to oligosphere differentiation. Linear sequence motif analyses of regulated class I phosphorylation sites.

**Table S6:** SILAC-based proximity biotinylation analysis for Dclk1 isoform interactomes. Interactome analysis based on protein level quantification, enrichment determined by relative quantification relative to heavy SILAC labeled control cells.

**Table S7:** SILAC-based proximity biotinylation analysis for Dclk1 isoform interactomes. Gene ontology (GO) analysis of individual protein level datasets.

**Table S8:** SILAC-based proximity biotinylation analysis for Dclk1 isoform interactomes. Differential expression analysis of individual protein level datasets.

**Table S9:** SILAC-based proximity biotinylation analysis for Dclk1 isoform interactomes. GO analyses of differentially expressed proteins based on binary comparisons of individual protein level datasets.

**Table S10:** SILAC-based proximity biotinylation analysis for Dclk1 isoform interactomes. Identified class I phosphosites, enrichment determined by relative quantification relative to heavy SILAC labeled control cells.

**Table S11:** SILAC-based proximity biotinylation analysis for Dclk1 isoform interactomes. GO analysis of individual class I phosphosite-based datasets.

**Table S12:** SILAC-based proximity biotinylation analysis for Dclk1 isoform interactomes. Differential expression analysis of phosphorylation sites determined for individual constructs.

**Table S13:** Investigation of potential Dclk1 cleavage sites. Semi-tryptic search results for Dclk1 BioID constructs and software-based prediction of potential Dclk1 cleavage sites by calpain.

**Table S14:** Primer sequences utilized for cloning and site directed mutagenesis experiments.

**Supplementary File:** Uncropped western blots and agarose gels.

## References

1. Azevedo, F.A.C., Carvalho, L.R.B., Grinberg, L.T., Farfel, J.M., Ferretti, R.E.L., Leite, R.E.P., Jacob Filho, W., Lent, R., and Herculano-Houzel, S. (2009). Equal numbers of neuronal and nonneuronal cells make the human brain an isometrically scaled-up primate brain. The Journal of comparative neurology 513, 532–541. 10.1002/cne.21974.

2. Callaway, E.M., Dong, H.W., Ecker, J.R., Hawrylycz, M.J., Huang, Z.J., Lein, E.S., Ngai, J., Osten, P., Ren, B., Tolias, A.S., et al. (2021). A multimodal cell census and atlas of the mammalian primary motor cortex. Nature 598, 86–102. 10.1038/s41586-021-03950-0.

3. Allen, N.J., and Lyons, D.A. (2018). Glia as architects of central nervous system formation and function. Science 362, 181–185. 10.1126/science.aat0473.

4. Schoor, C., Brocke-Ahmadinejad, N., Gieselmann, V., and Winter, D. (2019). Investigation of Oligodendrocyte Precursor Cell Differentiation by Quantitative Proteomics. Proteomics 19, 11, 1900057. 10.1002/pmic.201900057.

5. Dimou, L., Simon, C., Kirchhoff, F., Takebayashi, H., and Götz, M. (2008). Progeny of Olig2-expressing progenitors in the gray and white matter of the adult mouse cerebral cortex. The Journal of neuroscience 28, 10434–10442. 10.1523/jneurosci.2831-08.2008.

6. Filippi, M., Bar-Or, A., Piehl, F., Preziosa, P., Solari, A., Vukusic, S., and Rocca, M.A. (2018). Multiple sclerosis. Nature reviews. Disease primers 4, 43. 10.1038/s41572-018-0041-4.

7. van der Knaap, M.S., and Bugiani, M. (2017). Leukodystrophies: a proposed classification system based on pathological changes and pathogenetic mechanisms. Acta neuropathologica 134, 351–382. 10.1007/s00401-017-1739-1.

8. Goldman, S.A., and Kuypers, N.J. (2015). How to make an oligodendrocyte. Development 142, 3983–3995. 10.1242/dev.126409.

9. Chang, A., Tourtellotte, W.W., Rudick, R., and Trapp, B.D. (2002). Premyelinating oligodendrocytes in chronic lesions of multiple sclerosis. N. Engl. J. Med. 346, 165–173. 10.1056/NEJMoa010994.

10. Nishiyama, A., Komitova, M., Suzuki, R., and Zhu, X.Q. (2009). Polydendrocytes (NG2 cells): multifunctional cells with lineage plasticity. Nat. Rev. Neurosci. 10, 9–22. 10.1038/nrn2495.

11. Schoor, C., Brocke-Ahmadinejad, N., Gieselmann, V., and Winter, D. (2019). Investigation of Oligodendrocyte Precursor Cell Differentiation by Quantitative Proteomics. Proteomics 19, e1900057, Journal Article. 10.1002/pmic.201900057.

12. Chaerkady, R., Letzen, B., Renuse, S., Sahasrabuddhe, N.A., Kumar, P., All, A.H., Thakor, N.V., Delanghe, B., Gearhart, J.D., Pandey, A., and Kerr, C.L. (2011). Quantitative temporal proteomic analysis of human embryonic stem cell differentiation into oligodendrocyte progenitor cells. Proteomics 11, 4007–4020. 10.1002/pmic.201100107.

13. Letzen, B.S., Liu, C., Thakor, N.V., Gearhart, J.D., All, A.H., and Kerr, C.L. (2010). MicroRNA Expression Profiling of Oligodendrocyte Differentiation from Human Embryonic Stem Cells. PLoS One 5, 12, e10480. 10.1371/journal.pone.0010480.

14. Hu, J.G., Fu, S.L., Zhang, K.H., Li, Y., Yin, L., Lu, P.H., and Xu, X.M. (2004). Differential gene expression in neural stem cells and oligodendrocyte precursor cells: A cDNA microarray analysis. Journal of Neuroscience Research 78, 637–646. 10.1002/jnr.20317.

15. Chen, Y., Balasubramaniyan, V., Peng, J., Hurlock, E.C., Tallquist, M., Li, J., and Lu, Q.R. (2007). Isolation and culture of rat and mouse oligodendrocyte precursor cells. Nature protocols 2, 1044–1051. 10.1038/nprot.2007.149.

16. Salim, K., Kehoe, L., Minkoff, M.S., Bilsland, J.G., Munoz-Sanjuan, I., and Guest, P.C. (2006). Identification of differentiating neural progenitor cell markers using shotgun isobaric tagging mass spectrometry. Stem cells and development 15, 461–470. 10.1089/scd.2006.15.461.

17. Singh, S.A., Winter, D., Kirchner, M., Chauhan, R., Ahmed, S., Ozlu, N., Tzur, A., Steen, J.A., and Steen, H. (2014). Co-regulation proteomics reveals substrates and mechanisms of APC/C-dependent degradation. Embo J. 33, 385–399. 10.1002/embj.201385876.

18. Jarjour, A.A., Zhang, H., Bauer, N., Ffrench-Constant, C., and Williams, A. (2012). In vitro modeling of central nervous system myelination and remyelination. Glia 60, 1–12. 10.1002/glia.21231.

19. Conti, L., Pollard, S.M., Gorba, T., Reitano, E., Toselli, M., Biella, G., Sun, Y., Sanzone, S., Ying, Q.-L., Cattaneo, E., and Smith, A. (2005). Niche-independent symmetrical self-renewal of a mammalian tissue stem cell. PLoS biology 3, e283. 10.1371/journal.pbio.0030283.

20. Jung, M., Krämer, E., Grzenkowski, M., Tang, K., Blakemore, W., Aguzzi, A., Khazaie, K., Chlichlia, K., Blankenfeld, G.v., and Kettenmann, H. (1995). Lines of murine oligodendroglial precursor cells immortalized by an activated neu tyrosine kinase show distinct degrees of interaction with axons in vitro and in vivo. The European journal of neuroscience 7, 1245–1265.

21. Fisher, C.L., and Pei, G.K. (1997). Modification of a PCR-based site-directed mutagenesis method. Biotechniques 23, 570–574. 10.2144/97234bm01.

22. Winter, D., and Steen, H. (2011). Optimization of cell lysis and protein digestion protocols for the analysis of HeLa S3 cells by LC-MS/MS. Proteomics 11, 4726–4730. 10.1002/pmic.201100162.

23. Muller, T., and Winter, D. (2017). Systematic Evaluation of Protein Reduction and Alkylation Reveals Massive Unspecific Side Effects by Iodine-containing Reagents. Mol. Cell. Proteomics 16, 1173–1187. 10.1074/mcp.M116.064048.

24. Rappsilber, J., Ishihama, Y., and Mann, M. (2003). Stop and go extraction tips for matrix-assisted laser desorption/ionization, nanoelectrospray, and LC/MS sample pretreatment in proteomics. Analytical chemistry 75, 663–670. 10.1021/ac026117i.

25. Villen, J., and Gygi, S.P. (2008). The SCX/IMAC enrichment approach for global phosphorylation analysis by mass spectrometry. Nature Protocols 3, 1630–1638. 10.1038/nprot.2008.150.

26. Boersema, P.J., Raijmakers, R., Lemeer, S., Mohammed, S., and Heck, A.J.R. (2009). Multiplex peptide stable isotope dimethyl labeling for quantitative proteomics. Nature Protocols 4, 484–494. 10.1038/nprot.2009.21.

27. Søren S. Jensen, M.R.L. (2007). Evaluation of the impact of some experimental procedures on different phosphopeptide enrichment techniques. Rapid Communications in Mass Spectrometry 21, 3635–3645.

28. Dehghani, A., Godderz, M., and Winter, D. (2018). Tip-Based Fractionation of Batch-Enriched Phosphopeptides Facilitates Easy and Robust Phosphoproteome Analysis. J. Proteome Res. 17, 46–54. 10.1021/acsj.proteome.7b00256.

29. Winter, D., Seidler, J., Ziv, Y., Shiloh, Y., and Lehmann, W.D. (2009). Citrate boosts the performance of phosphopeptide analysis by UPLC-ESI-MS/MS. J. Proteome Res. 8, 418–424, Journal Article. 10.1021/pr800304n.

30. Ritchie, M.E., Phipson, B., Wu, D., Hu, Y.F., Law, C.W., Shi, W., and Smyth, G.K. (2015). limma powers differential expression analyses for RNA-sequencing and microarray studies. Nucleic Acids Research 43, 13, e47. 10.1093/nar/gkv007.

31. Valikangas, T., Suomi, T., and Elo, L.L. (2018). A systematic evaluation of normalization methods in quantitative label-free proteomics. Brief. Bioinform. 19, 1–11. 10.1093/bib/bbw095.

32. Benjamini, Y., and Hochberg, Y. (1995). Controlling the false discoveryx rate – a practical and powerful approach to multiple testing. J. R. Stat. Soc. Ser. B-Stat. Methodol. 57, 289–300. 10.1111/j.2517-6161.1995.tb02031.x.

33. Ginestet, C. (2011). ggplot2: Elegant Graphics for Data Analysis. Journal of the Royal Statistical Society Series a-Statistics in Society 174, 245–245. 10.1111/j.1467-985X.2010.00676_9.x.

34. Cox, J., and Mann, M. (2008). MaxQuant enables high peptide identification rates, individualized p.p.b.-range mass accuracies and proteome-wide protein quantification. Nature biotechnology 26, 1367–1372. 10.1038/nbt.1511.

35. Tyanova, S., Temu, T., Sinitcyn, P., Carlson, A., Hein, M.Y., Geiger, T., Mann, M., and Cox, J. (2016). The Perseus computational platform for comprehensive analysis of (prote)omics data. Nature methods 13, 731–740. 10.1038/nmeth.3901.

36. Wickham, H., Averick, M., Bryan, J., Chang, W., McGowan, L.D.A., François, R., Grolemund, G., Hayes, A., Henry, L., and Hester, J. (2019). Welcome to the Tidyverse. Journal of open source software 4, 1686.

37. Team, R.C. (2013). R: A language and environment for statistical computing.

38. Colaert, N., Helsens, K., Martens, L., Vandekerckhove, J., and Gevaert, K. (2009). Improved visualization of protein consensus sequences by iceLogo. Nature Methods 6, 786–787. 10.1038/nmeth1109-786.

39. Wu, T.Z., Hu, E.Q., Xu, S.B., Chen, M.J., Guo, P.F., Dai, Z.H., Feng, T.Z., Zhou, L., Tang, W.L., Zhan, L., et al. (2021). clusterProfiler 4.0: A universal enrichment tool for interpreting omics data. Innovation-Amsterdam 2, 11, 100141. 10.1016/j.xinn.2021.100141.

40. Willforss, J., Chawade, A., and Levander, F. (2019). NormalyzerDE: Online Tool for Improved Normalization of Omics Expression Data and High-Sensitivity Differential Expression Analysis. J. Proteome Res. 18, 732–740. 10.1021/acs.jproteome.8b00523.

41. Zhang, Y.W., Denham, J., and Thies, R.S. (2006). Oligodendrocyte progenitor cells derived from human embryonic stem cells express neurotrophic factors. Stem cells and development 15, 943– 952. 10.1089/scd.2006.15.943.

42. Kerr, C.L., Letzen, B.S., Hill, C.M., Agrawal, G., Thakor, N.V., Sterneckert, J.L., Gearhart, J.D., and All, A.H. (2010). Efficient differentiation of human embryonic stem cells into oligodendrocyte progenitors for application in a rat contusion model of spinal cord injury. The International journal of neuroscience 120, 305–313. 10.3109/00207450903585290.

43. Ward, J.H. (1963). Hierarchical Grouping to Optimize an Objective Function. Journal of the American Statistical Association 58, 236. 10.2307/2282967.

44. Alonso, S.B., Schulze-Steikow, M., and Calegari, F. (2014). Cell cycle activity of neural precursors in the diseased mammalian brain. Frontiers in Neuroscience 8, 39. 10.3389/fnins.2014.00039.

45. Zhao, X.H., He, X.L., Han, X.L., Yu, Y., Ye, F., Chen, Y., Hoang, T., Xu, X.M., Mi, Q.S., Xin, M., et al. (2010). MicroRNA-Mediated Control of Oligodendrocyte Differentiation. Neuron 65, 612–626. 10.1016/j.neuron.2010.02.018.

46. Urban, N., and Guillemot, F. (2014). Neurogenesis in the embryonic and adult brain: same regulators, different roles. Frontiers in Cellular Neuroscience 8, 396. 10.3389/fncel.2014.00396.

47. Ji, S., Doucette, J.R., and Nazarali, A.J. (2011). Sirt2 is a novel in vivo downstream target of Nkx2.2 and enhances oligodendroglial cell differentiation. Journal of molecular cell biology 3, 351–359. 10.1093/jmcb/mjr009.

48. Tretiakova, A., Steplewski, A., Johnson, E.M., Khalili, K., and Amini, S. (1999). Regulation of myelin basic protein gene transcription by Sp1 and Puralpha: Evidence for association of Sp1 and Puralpha in brain. Journal of cellular physiology 181, 160–168. 10.1002/(sici)1097-4652(199910)181:1&<160::Aid-jcp17>3.0.Co;2-h.

49. Bernstein, H.-G., Smalla, K.-H., Dürrschmidt, D., Keilhoff, G., Dobrowolny, H., Steiner, J., Schmitt, A., Kreutz, M.R., and Bogerts, B. (2012). Increased density of prohibitin-immunoreactive oligodendrocytes in the dorsolateral prefrontal white matter of subjects with schizophrenia suggests extraneuronal roles for the protein in the disease. Neuromolecular medicine 14, 270– 280. 10.1007/s12017-012-8185-y.

50. Gurok, U., Steinhoff, C., Lipkowitz, B., Ropers, H.H., Scharff, C., and Nuber, U.A. (2004). Gene expression changes in the course of neural progenitor cell differentiation. Journal of Neuroscience 24, 5982–6002. 10.1523/jneurosci.0809-04.2004.

51. Wang, H.H., Xiao, Z.T., Zheng, J.L., Wu, J., Hu, X.L., Yang, X.R., and Shen, Q. (2019). ZEB1 Represses Neural Differentiation and Cooperates with CTBP2 to Dynamically Regulate Cell Migration during Neocortex Development. Cell Reports 27, 2335–2353. 10.1016/j.celrep.2019.04.081.

52. Ligon, K.L., Fancy, S.P.J., Franklin, R.J.M., and Rowitch, D.H. (2006). Olig gene function in CNS development and disease. Glia 54, 1–10. 10.1002/glia.20273.

53. Fernandez, A., Northcott, P.A., Dalton, J., Fraga, C., Ellison, D., Angers, S., Taylor, M.D., and Kenney, A.M. (2009). YAP1 is amplified and up-regulated in hedgehog-associated medulloblastomas and mediates Sonic hedgehog-driven neural precursor proliferation. Genes & Development 23, 2729–2741. 10.1101/gad.1824509.

54. Xiao, L.Y., Dang, Y.N., Hu, B.J., Luo, L., Zhao, P.P., Wang, S.H., and Zhang, K. Overlapping functions of RBBP4 and RBBP7 in regulating cell proliferation and histone H3.3 deposition during mouse preimplantation development. Epigenetics. 10.1080/15592294.2021.1999006.

55. Hu, J.-G., Wang, Y.-X., Wang, H.-J., Bao, M.-S., Wang, Z.-H., Ge, X., Wang, F.-C., Zhou, J.-S., and Lü, H.-Z. (2012). PDGF-AA mediates B104CM-induced oligodendrocyte precursor cell differentiation of embryonic neural stem cells through Erk, PI3K, and p38 signaling. Journal of molecular neuroscience : MN 46, 644–653. 10.1007/s12031-011-9652-x.

56. Zhou, J., Tien, A.C., Alberta, J.A., Ficarro, S.B., Griveau, A., Sun, Y., Deshpande, J.S., Card, J.D., Morgan-Smith, M., Michowski, W., et al. (2017). A Sequentially Priming Phosphorylation Cascade Activates the Gliomagenic Transcription Factor Olig2. Cell Reports 18, 3167–3177. 10.1016/j.celrep.2017.03.003.

57. Liu, X.Y., Li, C.H., Li, J., Xie, L.S., Hong, Z., Zheng, K., Zhao, X.F., Yang, A.F., Xu, X.F., Tao, H.P., et al. (2022). EGF signaling promotes the lineage conversion of astrocytes into oligodendrocytes. Molecular Medicine 28, 50. 10.1186/s10020-022-00478-5.

58. Zhang, Z.H., Li, J.J., Wang, Q.J., Zhao, W.Q., Hong, J., Lou, S.J., and Xu, X.H. (2015). WNK1 is involved in Nogo66 inhibition of OPC differentiation. Molecular and Cellular Neuroscience 65, 135–142. 10.1016/j.mcn.2015.03.003.

59. Boersema, P.J., Raijmakers, R., Lemeer, S., Mohammed, S., and Heck, A.J. (2009). Multiplex peptide stable isotope dimethyl labeling for quantitative proteomics. Nature protocols 4, 484–494. 10.1038/nprot.2009.21.

60. Hornbeck, P.V., Zhang, B., Murray, B., Kornhauser, J.M., Latham, V., and Skrzypek, E. (2015). PhosphoSitePlus, 2014: mutations, PTMs and recalibrations. Nucleic acids research 43, D512–520. 10.1093/nar/gku1267.

61. Sock, E., and Wegner, M. (2021). Using the lineage determinants Olig2 and Sox10 to explore transcriptional regulation of oligodendrocyte development. Developmental Neurobiology 81, 892–901. 10.1002/dneu.22849.

62. 62. Wang, Y., Zhang, Y.H., Zhang, S., Kim, B.K., Hull, V.L., Xu, J., Prabhu, P., Gregory, M., Martinez-Cerdeno, V., Zhan, X.H., et al. (2021). PARP1-mediated PARylation activity is essential for oligodendroglial differentiation and CNS myelination. Cell Reports 37, 109695. 10.1016/j.celrep.2021.109695.

63. Dai, Z.M., Sun, S.H., Wang, C.Y., Huang, H., Hu, X.M., Zhang, Z.Y., Lu, Q.R., and Qiu, M.S. (2014). Stage-Specific Regulation of Oligodendrocyte Development by Wnt/beta-Catenin Signaling. Journal of Neuroscience 34, 8467–8473. 10.1523/jneurosci.0311-14.2014.

64. Neriec, N., and Percipalle, P. (2018). Sorting mRNA Molecules for Cytoplasmic Transport and Localization. Frontiers in Genetics 9, 510. 10.3389/fgene.2018.00510.

65. Genethliou, N., Panayiotou, E., Panayi, H., Orford, M., Mean, R., Lapathitis, G., Gill, H., Raoof, S., De Gasperi, R., Elder, G., et al. (2009). SOX1 links the function of neural patterning and Notch signalling in the ventral spinal cord during the neuron-glial fate switch. Biochemical and Biophysical Research Communications 390, 1114–1120. 10.1016/j.bbrc.2009.08.154.

66. Tian, Y.M., Lei, L., and Minden, A. (2011). A key role for Pak4 in proliferation and differentiation of neural progenitor cells. Developmental Biology 353, 206–216. 10.1016/j.ydbio.2011.02.026.

67. Nakano, I., Paucar, A.A., Bajpai, R., Dougherty, J.D., Zewail, A., Kelly, T.K., Kim, K.J., Ou, J., Groszer, M., Imura, T., et al. (2005). Maternal embryonic leucine zipper kinase (MELK) regulates multipotent neural progenitor proliferation. Journal of Cell Biology 170, 413–427. 10.1083/jcb.200412115.

68. Silverman, M.A., Benard, O., Jaaro, H., Rattner, A., Citri, Y., and Seger, R. (1999). CPG16, a novel protein serine/threonine kinase downstream of cAMP-dependent protein kinase. The Journal of biological chemistry 274, 2631–2636.

69. Shang, L.M., Kwon, Y.G., Nandy, S., Lawrence, D.S., and Edelman, A.M. (2003). Catalytic and regulatory domains of doublecortin kinase-1. Biochemistry 42, 2185–2194. 10.1021/bi026913i.

70. Burgess, H.A., and Reiner, O. (2001). Cleavage of doublecortin-like kinase by calpain releases an active kinase fragment from a microtubule anchorage domain. The Journal of biological chemistry 276, 36397–36403. 10.1074/jbc.M105153200.

71. Koizumi, H., Fujioka, H., Togashi, K., Thompson, J., Yates III, J.R., Gleeson, J.G., and Emoto, K. (2017). DCLK1 phosphorylates the microtubule-associated protein MAP7D1 to promote axon elongation in cortical neurons. Developmental Neurobiology 77, 493–510. 10.1002/dneu.22428.

72. O’Connell, M.R., Sarkar, S., Luthra, G.K., Okugawa, Y., Toiyama, Y., Gajjar, A.H., Qiu, S., Goel, A., and Singh, P. (2015). Epigenetic changes and alternate promoter usage by human colon cancers for expressing DCLK1-isoforms: Clinical Implications. Scientific reports 5, 14983. 10.1038/srep14983.

73. Burgess, H.A., and Reiner, O. (2002). Alternative splice variants of doublecortin-like kinase are differentially expressed and have different kinase activities. The Journal of biological chemistry 277, 17696–17705. 10.1074/jbc.M111981200.

74. Dijkmans, T.F., van Hooijdonk, L.W., Fitzsimons, C.P., and Vreugdenhil, E. (2010). The doublecortin gene family and disorders of neuronal structure. Central nervous system agents in medicinal chemistry 10, 32–46. 10.2174/187152410790780118.

75. Winter, D., Kugelstadt, D., Seidler, J., Kappes, B., and Lehmann, W.D. (2009). Protein phosphorylation influences proteolytic cleavage and kinase substrate properties exemplified by analysis of in vitro phosphorylated Plasmodium falciparum glideosome-associated protein 45 by nano-ultra performance liquid chromatography-tandem mass spectrometry. Anal. Biochem. 393, 41–47. 10.1016/j.ab.2009.06.022.

76. Roux, K.J., Kim, D.I., Raida, M., and Burke, B. (2012). A promiscuous biotin ligase fusion protein identifies proximal and interacting proteins in mammalian cells. The Journal of cell biology 196, 801–810. 10.1083/jcb.201112098.

77. Agulto, R.L., Rogers, M.M., Tan, T.C., Ramkumar, A., Downing, A.M., Bodin, H., Castro, J., Nowakowski, D.W., and Ori-McKenney, K.M. (2021). Autoregulatory control of microtubule binding in doublecortin-like kinase 1. eLife 10, 22, e60126. 10.7554/eLife.60126.

78. Shu, T., Tseng, H.-C., Sapir, T., Stern, P., Zhou, Y., Sanada, K., Fischer, A., Coquelle, F.M., Reiner, O., and Tsai, L.-H. (2006). Doublecortin-like kinase controls neurogenesis by regulating mitotic spindles and M phase progression. Neuron 49, 25–39. 10.1016/j.neuron.2005.10.039.

79. Liu, Y., Ferguson, F.M., Li, L.B., Kuljanin, M., Mills, C.E., Subramanian, K., Harshbarger, W., Gondi, S., Wang, J.H., Sorger, P.K., et al. (2020). Chemical Biology Toolkit for DCLK1 Reveals Connection to RNA Processing. Cell Chem. Biol. 27, 1229–1240. 10.1016/j.chembiol.2020.07.011.

80. Mobley, R.J., Raghu, D., Duke, L.D., Abell-Hart, K., Zawistowski, J.S., Lutz, K., Gomez, S.M., Roy, S., Homayouni, R., Johnson, G.L., and Abell, A.N. (2017). MAP3K4 Controls the Chromatin Modifier HDAC6 during Trophoblast Stem Cell Epithelial-to-Mesenchymal Transition. Cell Reports 18, 2387–2400. 10.1016/j.celrep.2017.02.030.

81. Rodor, J., Pan, Q., Blencowe, B.J., Eyras, E., and Cáceres, J.F. (2016). The RNA-binding profile of Acinus, a peripheral component of the exon junction complex, reveals its role in splicing regulation. RNA (New York, N.Y.) 22, 1411–1426. 10.1261/rna.057158.116.

82. Xu, S., Lai, S.K., Sim, D.Y., Ang, W.S.L., Li, H.Y., and Roca, X. (2022). SRRM2 organizes splicing condensates to regulate alternative splicing. Nucleic Acids Res 50, 8599–8614. 10.1093/nar/gkac669.

83. Sivan, G., Aviner, R., and Elroy-Stein, O. (2011). Mitotic modulation of translation elongation factor 1 leads to hindered tRNA delivery to ribosomes. J Biol Chem 286, 27927–27935. 10.1074/jbc.M111.255810.

84. Rikiyama, T., Curtis, J., Oikawa, M., Zimonjic, D.B., Popescu, N., Murphy, B.A., Wilson, M.A., and Johnson, A.C. (2003). GCF2: expression and molecular analysis of repression. Biochimica et biophysica acta 1629, 15–25. 10.1016/s0167-4781(03)00156-8.

85. Wang, J., Park, J.W., Drissi, H., Wang, X., and Xu, R.H. (2014). Epigenetic regulation of miR-302 by JMJD1C inhibits neural differentiation of human embryonic stem cells. J Biol Chem 289, 2384–2395. 10.1074/jbc.M113.535799.

86. Kanzaki, N., Ogita, H., Komura, H., Ozaki, M., Sakamoto, Y., Majima, T., Ijuin, T., Takenawa, T., and Takai, Y. (2008). Involvement of the nectin-afadin complex in PDGF-induced cell survival. Journal of cell science 121, 2008–2017. 10.1242/jcs.024620.

87. Bonucci, M., Kuperwasser, N., Barbe, S., Koka, V., de Villeneuve, D., Zhang, C., Srivastava, N., Jia, X., Stokes, M.P., Bienaimé, F., et al. (2020). mTOR and S6K1 drive polycystic kidney by the control of Afadin-dependent oriented cell division. Nature communications 11, 3200. 10.1038/s41467-020-16978-z.

88. Shinkai-Ouchi, F., Koyama, S., Ono, Y., Hata, S., Ojima, K., Shindo, M., duVerle, D., Ueno, M., Kitamura, F., Doi, N., et al. (2016). Predictions of Cleavability of Calpain Proteolysis by Quantitative Structure-Activity Relationship Analysis Using Newly Determined Cleavage Sites and Catalytic Efficiencies of an Oligopeptide Array. Mol. Cell. Proteomics 15, 1262–1280. 10.1074/mcp.M115.053413.

89. Goto-Silva, L., Martins, M., Murillo, J.R., Souza, L.R.Q., Vitoria, G., Oliveira, J.T., Nascimento, J.M., Loiola, E.C., Nogueira, F.C.S., Domont, G.B., et al. (2021). Quantitative profiling of axonal guidance proteins during the differentiation of human neurospheres. BBA-Proteins Proteomics 1869, 10, 140656. 10.1016/j.bbapap.2021.140656.

90. Hughes, S.M., Moussavi-Harami, F., Sauter, S.L., and Davidson, B.L. (2002). Viral-mediated gene transfer to mouse primary neural progenitor cells. Mol. Ther. 5, 16–24. 10.1006/mthe.2001.0512.

91. Cronin, J.C., Loftus, S.K., Baxter, L.L., Swatkoski, S., Gucek, M., and Pavan, W.J. (2018). Identification and functional analysis of SOX10 phosphorylation sites in melanoma. PLoS One 13, 18, e0190834. 10.1371/journal.pone.0190834.

92. Pauklin, S., and Vallier, L. (2013). The Cell-Cycle State of Stem Cells Determines Cell Fate Propensity. Cell 155, 135–147. 10.1016/j.cell.2013.08.031.

93. Fu, X.Q., Brown, K.J., Yap, C.C., Winckler, B., Jaiswal, J.K., and Liu, J.S. (2013). Doublecortin (Dcx) Family Proteins Regulate Filamentous Actin Structure in Developing Neurons. Journal of Neuroscience 33, 709–721. 10.1523/jneurosci.4603-12.2013.

94. Nakanishi, Y., Seno, H., Fukuoka, A., Ueo, T., Yamaga, Y., Maruno, T., Nakanishi, N., Kanda, K., Komekado, H., Kawada, M., et al. (2013). Dclk1 distinguishes between tumor and normal stem cells in the intestine. Nature genetics 45, 98–103. 10.1038/ng.2481.

95. Sureban, S.M., Madhoun, M.F., May, R., Qu, D., Ali, N., Fazili, J., Weygant, N., Chandrakesan, P., Ding, K., Lightfoot, S.A., and Houchen, C.W. (2015). Plasma DCLK1 is a marker of hepatocellular carcinoma (HCC): Targeting DCLK1 prevents HCC tumor xenograft growth via a microRNA-dependent mechanism. Oncotarget 6, 37200–37215. 10.18632/oncotarget.5808.

96. Weygant, N., Qu, D., May, R., Tierney, R.M., Berry, W.L., Zhao, L., Agarwal, S., Chandrakesan, P., Chinthalapally, H.R., Murphy, N.T., et al. (2015). DCLK1 is a broadly dysregulated target against epithelial-mesenchymal transition, focal adhesion, and stemness in clear cell renal carcinoma. Oncotarget 6, 2193–2205. 10.18632/oncotarget.3059.

97. Gregory, M.A., and Hann, S.R. (2000). c-Myc Proteolysis by the Ubiquitin-Proteasome Pathway: Stabilization of c-Myc in Burkitt’s Lymphoma Cells. Molecular and Cellular Biology 20, 2423– 2435. 10.1128/MCB.20.7.2423-2435.2000.

98. Rogers, S., Wells, R., and Rechsteiner, M. (1986). Amino acid sequences common to rapidly degraded proteins: the PEST hypothesis. Science 234, 364–368. 10.1126/science.2876518.

99. Domina, A.M., Vrana, J.A., Gregory, M.A., Hann, S.R., and Craig, R.W. (2004). MCL1 is phosphorylated in the PEST region and stabilized upon ERK activation in viable cells, and at additional sites with cytotoxic okadaic acid or taxol. Oncogene 23, 5301–5315. 10.1038/sj.onc.1207692.

100. Ariake, K., Ohtsuka, H., Motoi, F., Douchi, D., Oikawa, M., Rikiyama, T., Fukase, K., Katayose, Y., Egawa, S., and Unno, M. (2012). GCF2/LRRFIP1 promotes colorectal cancer metastasis and liver invasion through integrin-dependent RhoA activation. Cancer Lett. 325, 99–107. 10.1016/j.canlet.2012.06.012.

101. Perez-Riverol, Y., Csordas, A., Bai, J.W., Bernal-Llinares, M., Hewapathirana, S., Kundu, D.J., Inuganti, A., Griss, J., Mayer, G., Eisenacher, M., et al. (2019). The PRIDE database and related tools and resources in 2019: improving support for quantification data. Nucleic Acids Research 47, D442–D450. 10.1093/nar/gky1106.

